# Proteo-transcriptomics and morphometrics of teleost cardiac cells define regulatory networks and exercise-induced cardiomyocyte hypertrophy and hyperplasia

**DOI:** 10.64898/2026.01.11.698907

**Authors:** Osvaldo Contreras, Gabrielle D. Smith, Celine Santiago, Moumitha Dey, Chris Thekkedam, Renee Chand, Álvaro González-Rajal, Ling Zhong, John G. Lock, Emily Wong, Diane Fatkin, Richard P. Harvey

**Author notes:** These authors contributed equally (Gabrielle D. Smith, Celine Santiago, Moumitha Dey). **Correspondence:** Dr. Osvaldo Contreras, MSc, PhD Developmental and Regenerative Biology Division Victor Chang Cardiac Research Institute Lowy Packer Building, Level 7 405 Liverpool Street Darlinghurst, NSW 2010, Australia.

## Abstract

Zebrafish and medaka are powerful cardiovascular models, yet cellular and molecular investigations of adult heart cells have been constrained by suboptimal dissociation and characterization methods. To overcome these barriers, we developed a physiological-temperature workflow that generates high-yield, viable single-cell suspensions for FACS, imaging, and low-input molecular profiling. Using transgenic fluorescent reporters, we consistently isolate ∼6,000 cardiomyocytes per adult zebrafish ventricle and ∼12,000 from medaka, preserving cellular, structural, and molecular integrity. Single-cell morphometrics revealed cardiomyocyte heterogeneity and demonstrated that swimming exercise induces both hypertrophy and hyperplasia, while ventricular injury triggers expansion of regenerative *gata4*⁺ cardiomyocytes. We integrated proteomics and RNA-seq from FACS-purified cells to construct cell-type-specific proteo-transcriptomic atlases. Functional enrichment, transcription factor, and network analyses identified protein hubs and regulatory circuits defining cardiomyocyte and endothelial cell identity. Our platform delivers cell-type-resolved molecular datasets of adult teleost cardiac cells, establishing a systems-level resource for heart regeneration research, cardiovascular disease modeling, and drug discovery.

**Teaser:** A robust workflow for isolation, FACS, imaging, and multi-omics profiling of cardiomyocytes and cardiac endothelial cells in fish.

**HIGHLIGHTS:**

- Temperature-optimized dissociation and standardized FACS enable high-yield isolation of viable cardiac cells.
- Single-cell imaging uncovers morphological heterogeneity in adult ventricular cardiomyocytes.
- Sustained exercise induces both hypertrophy and hyperplasia of zebrafish cardiomyocytes.
- Proteo-transcriptomics defines core molecular programs and interaction networks in cardiomyocytes and endothelial cells.

## INTRODUCTION

Teleost fish, particularly zebrafish (*Danio rerio*) and medaka (*Oryzias latipes*), have emerged as invaluable ectothermic vertebrate models for cardiovascular research due to their unique traits: genetic tractability, optical transparency during development, and remarkable cardiac regenerative capacity. Unlike mammals, adult zebrafish are capable of fully regenerating their hearts following apical resection^1–4^, making them ideal for investigating the evolutionary cellular and molecular mechanisms underlying cardiac repair and regeneration^5^. In contrast, medaka cannot fully regenerate their hearts, but retain latent regenerative capacity^6–8^. These differences are attributed, in part, to divergent immune, endocardial/epicardial, and cardiomyocyte-specific responses^8–12^. Zebrafish cardiac regenerative prowess is largely driven by the dedifferentiation and proliferation of cardiomyocytes, including dedifferentiated cardiomyocytes that express the transcription factor gata4^1,2,5,11,13–15^—a process that has captured the attention of researchers seeking to unlock therapeutic strategies for human heart disease^5,10,16^.

The accessibility of transgenic zebrafish lines expressing fluorescent reporters in specific cardiac cell populations has further enhanced their utility for research^17,18^. As of November 2025, the Zebrafish Information Network (ZFIN)^19^ displays 10,657 transgenic zebrafish lines expressing GFP, including powerful tools such as *Tg(cmlc2:GFP)*^20,21^, which labels cardiomyocytes, and *Tg(kdrl/flk1:DsRed2)*^10^, which marks endothelial/endocardial cells, enabling visualization, tracking, targeting and isolation of distinct cardiac cell types for downstream analyses. These genetic resources, combined with key developmental and physiological similarities to mammalian cardiovascular systems and substantial protein-coding gene conservation between teleosts and humans^22^, position teleost models at the forefront of cardiac electrophysiology and cardiovascular disease research^23–25^.

Beyond regeneration and disease modeling, zebrafish have emerged as valuable tools for studying exercise-induced cardiac adaptations. While studies in humans and rodents demonstrate that regular physical activity enhances cardiac performance and systemic cardiometabolic health^26–28^, the cellular mechanisms underlying these benefits remain incompletely understood. Zebrafish offer a unique opportunity to dissect these mechanisms at cellular and molecular resolution. While foundational studies explored skeletal muscle plasticity using swimming regimens^29,30^, cardiac-focused studies demonstrated that swimming exercise is a powerful stimulus for cardiac plasticity^31–35^. In healthy zebrafish hearts, exercise induced significant cardiomyocyte proliferation (EdU⁺/Mef2⁺ cells) without altering overall cardiac function, and this capacity became critical during injury recovery, where exercised hearts showed a remarkable 4-fold increase in cardiomyocyte proliferation over non-exercised injured fish^31^. Another exercise study reported whole-heart enlargement, increased ventricular cross-sectional area, and thickened compact myocardium^33^. However, these zebrafish studies relied primarily on histology and bulk tissue assessments and the impact of exercise on individual cardiomyocyte morphology and the extent of hypertrophy at single-cell resolution remains unclear.

Despite the widespread adoption of zebrafish and, increasingly medaka, in cardiovascular research^7,24^ and comparative biology^8,9,36,37^, significant technical barriers have limited the full exploitation of these models. Current protocols for dissociating adult cardiac tissue into viable single-cell suspensions face several critical limitations, hindering comprehensive cellular and molecular analyses.

First, existing dissociation methods often employ enzymatic conditions optimized for mammalian systems^13,38–44^, typically utilizing temperatures of 32–37°C that exceed the physiological range of teleost fish (25–28°C)^45^. This temperature mismatch can induce thermal and cellular stress, compromise cell viability, and potentially introduce transcriptional artifacts that confound downstream analyses due to medium/low RNA quality as determined by the RNA Integrity Number (RIN)^38^. The resulting cell death and structural damage, often unreported in published protocols and articles^13,39–41,46^, limit both the yield and quality of isolated cardiac cells.

Second, the field lacks standardized, comprehensive protocols for flow cytometric characterization and fluorescence-activated cell sorting (FACS) of cardiac cell populations^39^. Many published studies provide insufficient detail regarding flow cytometry gating strategies, yield quantification, morphological determination, or quality assessment of isolated cells^38–40,47^. These methodological gaps have created reproducibility challenges, limited cross-laboratory comparisons, and led to the adoption of non-standardized approaches across institutions. Third, several existing protocols require extensive processing times, often exceeding several hours, which can further compromise cell viability and introduce additional variables that may affect experimental outcomes^40^. Moreover, the use of 70µm cell strainers and prolonged dissociation procedures^44^ may limit the throughput necessary for large-scale studies or time-sensitive downstream applications.

The development of robust, high-yield cardiac cell isolation protocols is essential for advancing several critical areas of cardiovascular research. Detailed morphological and functional characterization of cardiac cells necessitates the preservation of cellular architecture, as well as physiological and molecular properties. Harsh dissociation conditions can disrupt sarcomeric organization in cardiomyocytes, alter membrane integrity in cardiac stromal cells (e.g., endothelial cells and fibroblasts), and compromise the very features researchers seek to study. Moreover, the growing interest in cardiac regeneration has highlighted the need for methods capable of isolating and characterizing rare cell populations, such as dedifferentiated gata4⁺ cardiomyocytes^2,47^. These populations are often present in low numbers, requiring highly sensitive, rapid, accurate, and specific isolation techniques for meaningful analysis.

To address these technological and knowledge gaps, we developed and validated a fast, temperature-optimized enzymatic dissociation protocol capable of generating high-yield, viable single-cell suspensions from transgenic adult zebrafish and medaka hearts, and established standardized flow cytometry gating strategies and FACS protocols for robust identification and isolation of distinct cardiac cell populations. We demonstrated that optimizing dissociation conditions to match the physiological temperature of teleost fish significantly improved cell yield and viability while preserving cellular structure, function, and molecular profiles. Through microscopy and morphometrics, we provided comprehensive morphological characterization of isolated cardiomyocytes at baseline and in response to exercise or resection-induced cardiac remodeling, revealing an unexpected heterogeneity in cardiomyocytes morphologies. Finally, we leveraged our optimized cell isolation workflow to perform high-quality, low-input transcriptomic and proteomic quantitative mapping of ventricular cardiomyocytes and endothelial cells. Our analyses revealed cell-type-specific protein-protein interaction networks, transcription factor regulomes, gene regulatory networks (GRNs), and functional enrichments that define cell identities, establishing a blueprint framework that can be extended to other cell types and disease states.

## RESULTS

### A Fast and Optimized Cardiomyocyte Isolation and Characterization Protocol for Teleost Hearts

To better mimic zebrafish physiology, minimize temperature-induced artifacts, and optimize enzymatic efficiency while reducing processing time, we investigated whether a novel in-house enzymatic combination^48^ could enable efficient dissociation at lower temperatures and facilitate the isolation of high-quality cardiomyocytes. We developed and optimized a custom dissociation protocol employing papain (a highly efficient cysteine protease from papaya latex), Collagenase type II, Dispase, and DNase I. Given that these enzymes retain substantial activity below human body temperature, we directly compared compared efficiency and cell quality at 37°C and 28°C using ventricular cardiac tissue (**Fig. 1A**). Our optimized dissociation solution and protocol proved successful at 28°C, enabling rapid and efficient dissociation of ventricular tissue into single cells within 30–45 minutes. Compared to the conventional enzyme protocol at 37°C, dissociation at 28°C yielded substantially lower cell death rates (under 5% at 28°C, versus approximately 20% at 37°C; difference between means [37°C–28°C] ± SEM = 15.33±2.97; **Fig. 1B**). The protocol was equally effective for isolating adult ventricular cardiomyocytes from two transgenic *cmlc2:EGFP* teleost species (zebrafish and medaka) of similar body size, yielding comparable proportions of GFP-positive events across species (**Fig. 1C**). Flow cytometric analysis further confirmed that the majority of dissociated cells formed viable, single-cell suspensions, enabling reliable isolation of large numbers of live, GFP⁺ ventricular cardiomyocytes (**Fig. S1**).

**Fig. 1.**
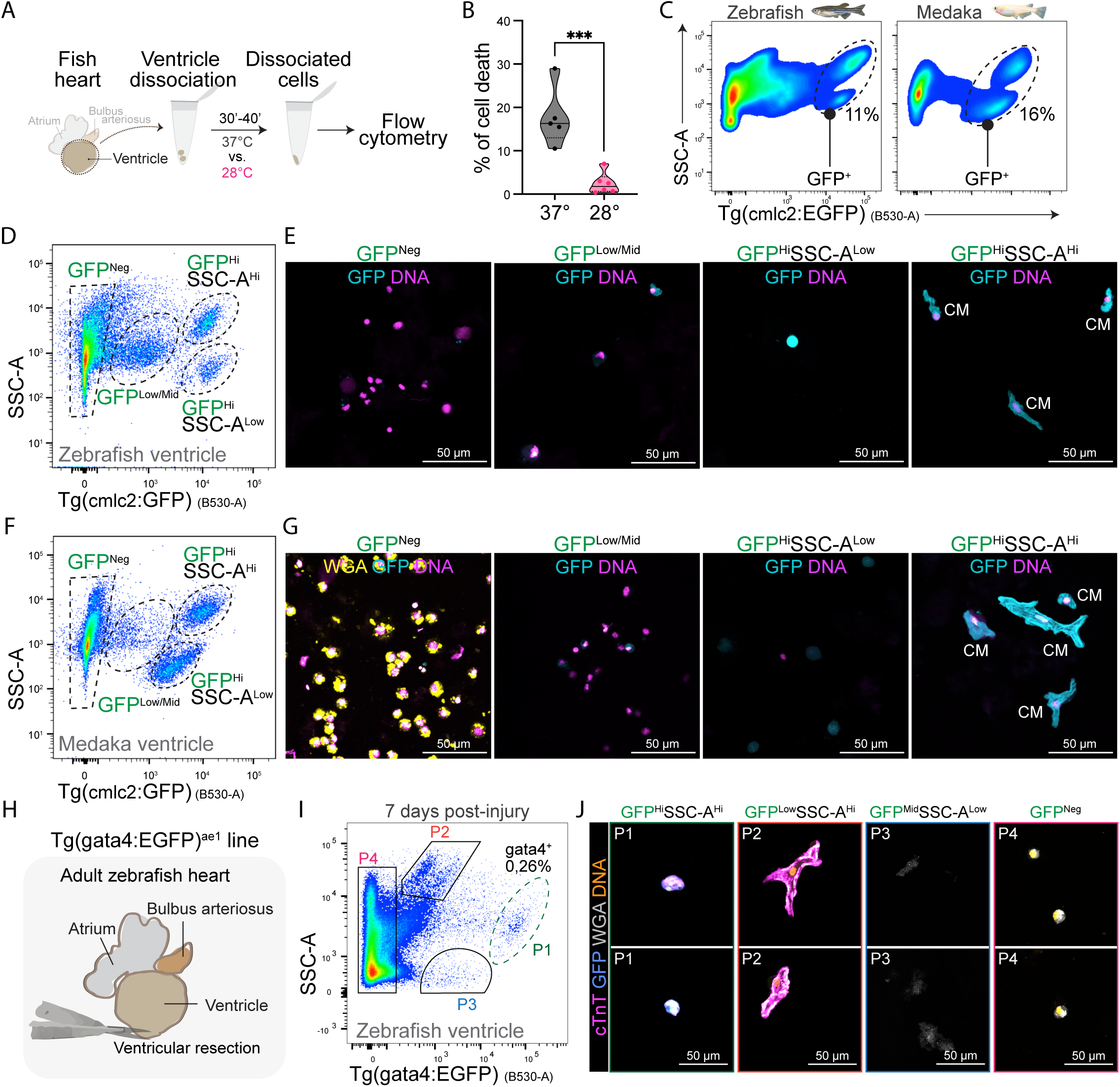
Dissociation and characterization of ventricular cardiomyocytes in zebrafish and medaka via fluorescence-activated cell sorting and imaging. (A) Outline of fish ventricle isolation, dissociation, and flow cytometry, comparing two dissociation temperatures using adult male fish. (B) Quantification of the percentage of dead cells under two distinct dissociation temperatures (°C). ***P<0.001 by nonparametric Mann-Whitney test comparing ranks; n=5-6, totalling 18 male ventricles. (C) Post-dissociation flow cytometry determination of GFP-labelled cardiomyocytes in combination with SSC-A, showing representative plots in two different fish species. (n = 6) (D) Post-dissociation flow cytometry analysis of zebrafish ventricular live cells, displaying four distinct clusters based on SSC-A and GFP fluorescence intensity. (E) Representative laser confocal images of the four primary flow cytometry clusters identified in (A). GFP fluorescence is shown in cyan, whereas DNA staining is shown in magenta hot. The GFP^Hi^SSC-A^Hi^ cluster is the onlyone containing viable and high-quality cardiomyocytes. (F) Post-dissociation flow cytometry analysis of medaka ventricular live cells, showing four distinct clusters based on SSC-A and GFP fluorescence intensity. (G) Representative laser confocal images of the four primary flow cytometry clusters identified in (C). GFP fluorescence is shown in cyan, and nuclei in magenta hot. The GFP^Hi^SSC-A^Hi^ clusteris the only one containing viable and high-quality cardiomyocytes. WGA (yellow) staining in the GFP^Neg^ panel indicates intact cell membranes in most GFP^Neg^ cells. CM, cardiomyocyte. (H) Outline of ventricular resection using the Tg(gata4:EGFP)^ae1^ reporter line. (I) Representative flow cytometry gating strategy to characterize distinct populations (P) based on SSC-A and GFP fluorescence intensity. (J) Representative laser confocal images of the four primary flow cytometry clusters (P1-P4) identified in (F) showing P1 (green dashed line) as the true gata4+ cells.

### Refining Flow Cytometric and FACS-Assisted Characterization of Adult *cmlc2:EGFP* Cardiomyocytes

Despite increasing efforts to characterize dissociated zebrafish cardiac cells, particularly adult cardiomyocytes, two critical aspects remain insufficiently addressed in the literature: (i) the establishment of comprehensive flow cytometry gating strategies with representative plots of pure cardiomyocytes^38–40^, and (ii) quantitative reporting of the quality and yield of isolated cardiac cell populations^39,40,49^. To address these gaps, we established a comprehensive flow cytometry-based evaluation of GFP⁺ ventricular cardiomyocytes using male *cmlc2:EGFP* adult zebrafish and medaka.

Our initial analysis of single-cell GFP⁺ events revealed two major cell clusters distinguished by side scatter characteristics (SSC-A^high^ versus SSC-A^low^), both exhibiting relatively high GFP fluorescence intensity (**Figure 1C** and **Fig. S1**). To determine which SSC-A cluster accurately represents true GFP⁺ ventricular cardiomyocytes, we combined FACS with cytospin-coupled confocal imaging. Detailed flow cytometric profiling identified two distinct GFP⁺ populations: GFP^high^/SSC-A^high^ and GFP^high^/SSC-A^low^(**Fig. 1D**). We hypothesized that the GFP^high^/SSC-A^high^ population consisted of intact cardiomyocytes due to its higher side scatter properties. FACS-assisted confocal microscopy confirmed this, showing that GFP^high^/SSC-A^high^ cells were viable, single-cell cardiomyocytes (**Fig. 1E**), whereas the GFP^high^/SSC-A^low^ population primarily consisted of cardiomyocyte debris, likely generated during tissue dissociation (**Fig. 1E**).

We further characterized additional flow cytometry populations. The GFP-negative fraction comprised non-cardiomyocyte cells, while GFP^low^/^mid^ populations likely represented either non-cardiomyocytes or remnants of structurally compromised GFP⁺ cardiomyocytes (**Fig. 1D, E**). Similar results in adult medaka ventricular tissue (**Fig. 1F, G**) reinforced the robustness and cross-species applicability of our dissociation and flow cytometry strategies. These findings were further validated using *cmlc2:EGFP;cmlc2:DsRed2-nuc* double-transgenic zebrafish, which helped mitigate potential confounding from autofluorescence in the GFP channel and provided orthogonal, compartment-resolved confirmation of cardiomyocyte identity (**Fig. S2A–C**). Our refined flow cytometry strategy also proved effective following PFA fixation, enabling clear distinction of nucleated (DNA⁺) GFP^high^ cardiomyocytes from nucleated GFP⁻ cells and debris (**Fig. S2D, E**). As expected, fixed cardiomyocytes (DNA⁺/GFP^high^) exhibited higher FSC-A values than DNA⁺/GFP⁻ cells (**Fig. S2D**). Thus, this comprehensive cytometric and imaging-based approach identifies the GFP^high^/SSC-A^high^ population as a definitive marker of viable, dissociated cardiomyocytes.

To complement our whole-cell analysis, we extended our approach to the nuclear level by isolating and characterizing single nuclei from dissociated adult ventricular tissue (**Fig. S3**). Flow cytometric analysis enabled identification of DNA-stained nuclei by side scatter area (SSC-A), allowing clear discrimination of nuclear populations (**Fig. S3A**). Comparative flow cytometry of single nuclei from adult zebrafish and medaka ventricles revealed distinct DNA fluorescence intensity profiles (**Fig. S3B, C**), suggesting species-specific differences in nuclear content or chromatin organization. Confocal microscopy confirmed these events as nuclei (**Fig. S3B, C**). This nuclear-level analysis provides an additional layer of validation for our cardiomyocyte identification strategy and offers a robust platform for future studies investigating nuclear dynamics, single-nucleus omics, ploidy, and transcriptional or epigenomic states in cardiac cells across species.

### Embryonic Profiling of Cardiomyocytes Using Flow Cytometry

To further validate our flow cytometry-based strategy and assess its applicability across developmental stages, we extended our dissociation analysis to embryonic zebrafish cardiomyocytes using the *cmlc2:EGFP* line (**Fig. S4**). After whole-embryo dissociation, single-cell suspensions from 28.5 hpf embryos were subjected to flow cytometry, and live cells were gated based on Zombie Yellow⁻/DAPI⁻ exclusion (**Fig. S4A**). Within this live population, GFP⁺ cardiomyocytes were clearly identifiable, demonstrating the feasibility of our gating strategy in early developmental contexts.

To estimate cell size, we correlated forward scatter area (FSC-A) with the diameter of calibration beads of known sizes (**Fig. S4B**)^50^. This calibration enabled quantification of embryonic GFP⁺ cardiomyocyte size distribution (**Fig. S4C**), providing a benchmark for interpreting FSC-A values in adult cardiomyocyte populations.

### Evaluation of Dedifferentiated *gata4⁺* Cardiomyocytes Post-Ventricular Resection

The adult zebrafish heart exhibits remarkable regenerative capacity, primarily driven by the dedifferentiation of mature cardiomyocytes and the activation of *gata4*-expressing progenitor-like cells following injury, as well as subsequent cardiomyocyte re-differentiation^1,2,51^. This complex regenerative response highlights the need for precise profiling of *gata4⁺* cardiomyocytes following cardiac damage^47^. To address this, we extended our dissociation-based flow cytometry workflow and performed a proof-of-concept analysis using the *Tg(-14.8gata4:GFP)^ae^*^1^ zebrafish line^52^ at 7 days post-ventricular resection (**Fig. 1H**).

At 7 days post-injury (7dpi), flow cytometry resolved four distinct single-cell clusters (P1–P4) (**Fig. 1I, Fig. S5**). Each cluster was characterized by FACS-assisted cytospin followed by confocal microscopy (**Fig. 1J**). Through co-staining for cardiac troponin T (TNNT2/cTnT), wheat germ agglutinin (WGA), and nuclear DNA—together with endogenous GFP fluorescence—we identified cluster P1 as dedifferentiating *gata4⁺* cardiomyocytes (**Fig. 1J**). They exhibited a rounded morphology consistent with the dedifferentiated, progenitor-like state. Notably, *gata4⁺* cells expanded by approximately three-to fourfold relative to uninjured and sham-operated controls (**Fig. S5C, D**). Interestingly, thoracotomy (sham) also increased the number of *gata4⁺* cells, potentially through preconditioning that activates regenerative programs^53^ or via sham-induced epicardial inflammation, which alone can trigger cardiomyocyte proliferation^3^.

The P2 cluster consisted of non-regenerative, *gata4⁻* cardiomyocytes, which displayed elevated autofluorescence in the GFP channel (B530-A) (**Fig. 1J**), likely reflecting injury-induced metabolic changes, high contractile protein abundance, and their large size. The P3 cluster was characterised by low side scatter (SSC-A^low^) and low GFP intensity, consistent with cellular debris (**Fig. 1J**). In contrast, the P4 cluster comprised non-cardiomyocyte cells with intact WGA-labelled membranes and preserved nuclear and cellular structure (**Fig. 1J**). Together, these findings validate our dissociation approach and FACS-assisted imaging workflows for profiling low-abundance *gata4⁺* regenerating cardiomyocytes.

### Ventricular Cardiomyocyte Yields Across the Adult Zebrafish Lifespan

No comprehensive reports have systematically evaluated ventricular cardiomyocyte yields in zebrafish or medaka, hindering accurate calculation of the number of fish or heart/ventricle mass required for experiments. In response, we quantified cardiomyocyte yields from male zebrafish ventricles at three adult stages using *cmlc2:EGFP* fish: young adults (4–5 months), mature adults (6–10 months), and aged individuals (12–16 months) (**Fig. 2A**). Cardiomyocyte counts were normalized to both resuspension volume and animal weight (grams) to ensure consistency (**Fig. 2A, B**). Flow cytometry revealed that young and mature adults consistently yielded approximately 4,000–7,000 GFP⁺ ventricular cardiomyocytes per ventricle, with an average of ∼6,000 live cardiomyocytes in adults (**Fig. 2B, C**). In contrast, aged zebrafish exhibited a significant decline, averaging around 2,000–3,000 GFP⁺ cells per ventricle, indicating an age-dependent reduction in recoverable cardiomyocyte numbers (**Fig. 2B, C**). This pattern was consistent across different fish cohorts and experimental days.

**Fig. 2.**
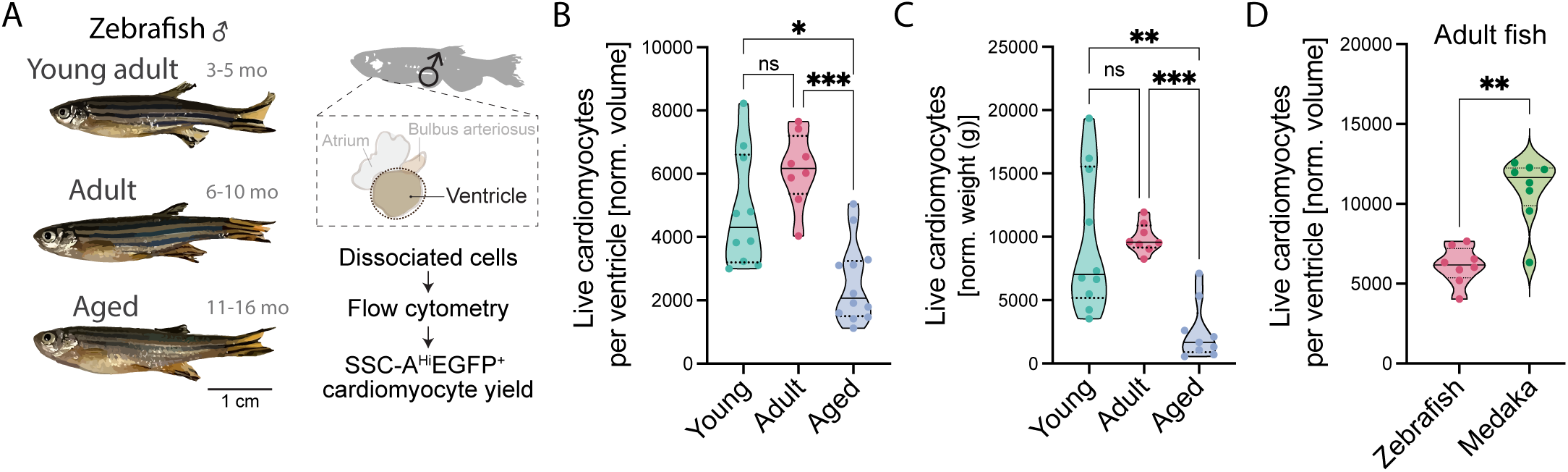
Quantification of ventricular cardiomyocyte yield across the adult zebrafish lifespan and comparison with medaka. (A) Representative adult zebrafish illustrations showing relative size of three adult types: young adult, adult, and aged male zebrafish, later used to evaluate cardiomyocyte yield, and outline of cell isolation and yield evaluation of SSC-A^Hi^/EGFP^Hi^ ventricu-lar cardiomyocytes from adult male zebrafish. (B) Cardiomyocyte yield per ventricle comparing young adult, adult, and aged male fish, normalized by final total volume per dissociaton. ***P<0.0005, *P<0.05 by one-way ANOVA with Kruskal-Wallis’s post-test; n=8-10. (C) Cardiomyocyte yield per ventricle comparing young adult, adult, and aged male fish, normalized to body weight. ***P = 0.0007; **P<0.005 by one-way ANOVA with Kruskal-Wallis post-test and Dunn’s multiple comparisons; n=8-10. (D) Yield of live ventricular cardiomyocytes per ventricle in adult (6-10-month-old) male zebrafish versus medaka, normalized by final dissociation volume, revealing significantly higher yields in medaka. **P<0.005 by nonparametric Mann-Whitney test comparing ranks; n=8, totalling 24 male ventricles.

To evaluate cross-species applicability and benchmark yield, we also compared ventricular cardiomyocyte numbers between adult zebrafish and medaka. Notably, male medaka ventricles yielded significantly more live GFP^Hi^/SSC-A^Hi^ cardiomyocytes per ventricle than zebrafish (median: 11,646 vs. 6,174; actual median difference: 5,472; P = 0.0012, Mann-Whitney test; **Fig. 2D**), further supporting cross-species robustness and high efficiency of our protocol.

### Morphological Assessment and Sarcomeric Staining of Pure Adult Zebrafish Ventricular Cardiomyocytes

To further validate the structural integrity of adult zebrafish cardiomyocytes isolated using our enzymatic dissociation protocol, we obtained pure adult ventricular cardiomyocytes from the *cmlc2:EGFP* line via FACS and performed immunostaining for key sarcomeric proteins (**Fig. 3A**). Given that sarcomeres are the fundamental contractile units of cardiomyocytes, their organization serves as a reliable indicator of overall cellular structure.

**Fig. 3.**
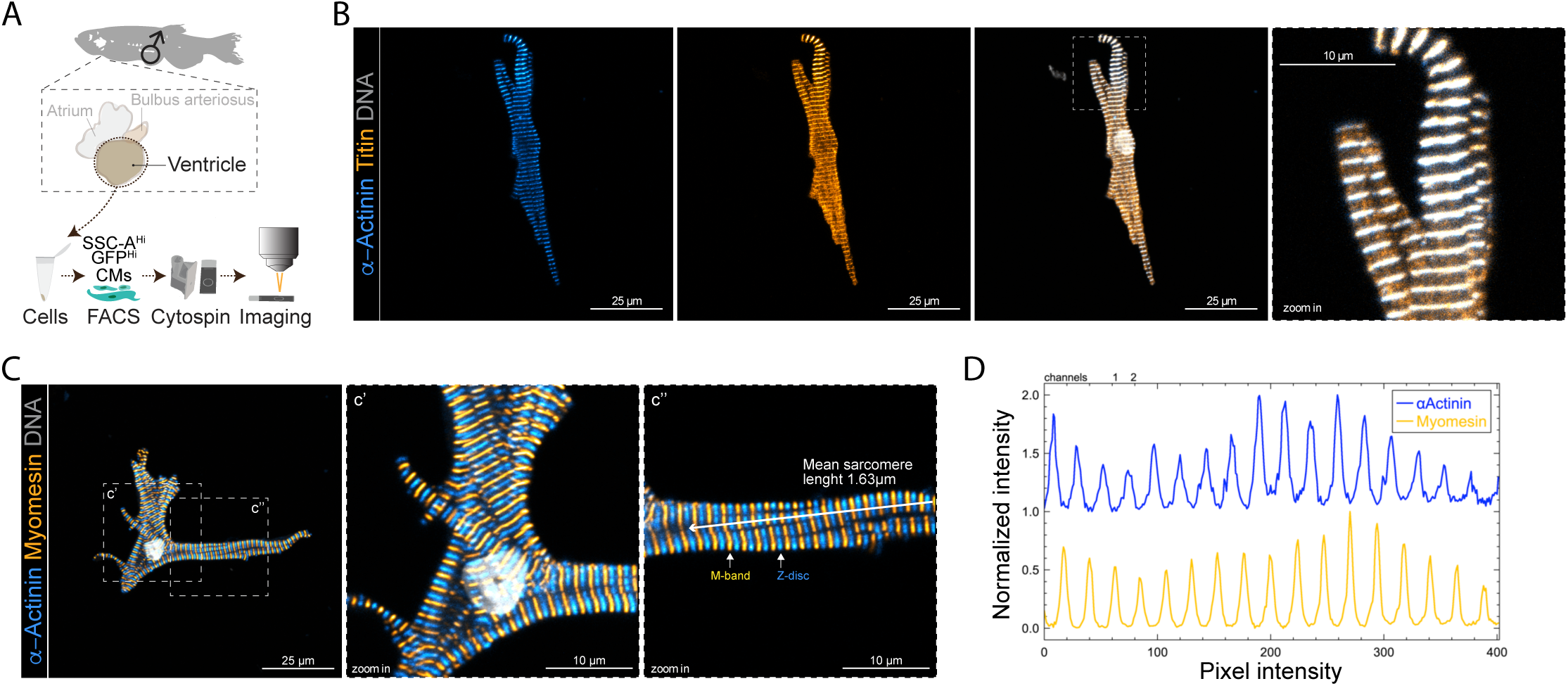
FACS-assisted characterization and staining of adult ventricular zebrafish cardiomyocytes. (A) Outline of cell isolation and FACS of SSC-A^Hi^/GFP^Hi^ ventricular cardiomyocytes from adult male zebrafish. (B) Representative laser confocal z-stack images of a sorted cardiomyocyte showing the colocalization of two sarcomeric proteins. α-Actinin staining is shown in cyan hot, Titin in orange hot, and nuclear staining in grey. The image on the right shows a magnified (3x zoom) area. (C) Representative laser confocal z-stack images of a sorted cardiomyocyte showing the sarcomeric localization of α-Actinin and myomesin. α-Actinin staining is shown in cyan hot, Myomesin in orange hot, and nuclear staining in grey. The images on the right (c’ and c’’) show 3x magnified regions highlighting the M-bands and Z-discs. (D) Normalized fluorescence intensity profile showing sarcomeric α-Actinin and Myomesin staining along the depicted line shown in c’’. The M-bands are labeled by Myomesin and the Z-discs by α-Actinin.

FACS-isolated adult ventricular cardiomyocytes were successfully stained for sarcomeric α-Actinin and Titin, which revealed well-preserved, regularly spaced Z-discs^54^ (**Fig. 3B**). DNA dye staining remained intact, confirming preservation of nuclear structure. Co-labeling with α-Actinin and Myomesin (an M-line protein) further validated the integrity of both Z-discs and M-lines, with confocal imaging revealing distinct sarcomeric and myofibrillar banding patterns across diverse cardiomyocyte morphologies (**Fig. 3C**). Thin and thick filaments were organized into myofibrils aligned along the main axes of the cardiomyocytes, displaying regularly spaced Z-discs. Notably, substantial myofibril branching was also observed (**Fig. 3C**). These sarcomeric markers enabled fluorescence intensity profiling along parallel myofibrils, allowing precise quantification of sarcomere length at approximately 1.6 μm (**Fig. 3C, D**), consistent with previous reports^5^.

Because we observed complex ventricular cardiomyocyte morphologies distinct from the "elongated" phenotype described by others^49,55^, we further explored cardiomyocyte shapes using cytospun cells immediately after dissociation and PFA fixation (**Fig. 4A**). We found that the majority of cytospun GFP⁺ ventricular cardiomyocytes exhibited excellent cellular and sarcomeric integrity, with distinct intracellular myofibrils (**Fig. 4B**). A broad range of morphologies—including rounded, rod-shaped, multipolar, and elongated cells—was observed (**Fig. 4C**). To expand this analysis beyond GFP-based imaging, we performed α-Actinin and Myomesin staining and used tiling confocal microscopy to capture morphological diversity across a wider field of view (**Fig. 4D**). These morphologies were identified in intact, live, nucleated *cmlc2*⁺ cardiomyocytes isolated by FACS, with α-Actinin and Myomesin immunostaining facilitating detailed visualization of sarcomeric organization within individual myofibrils (**Fig. 4D, E**). **Figure 4F** further illustrates this heterogeneity, with notable prevalence of rounded and rod-shaped cardiomyocytes.

**Fig. 4.**
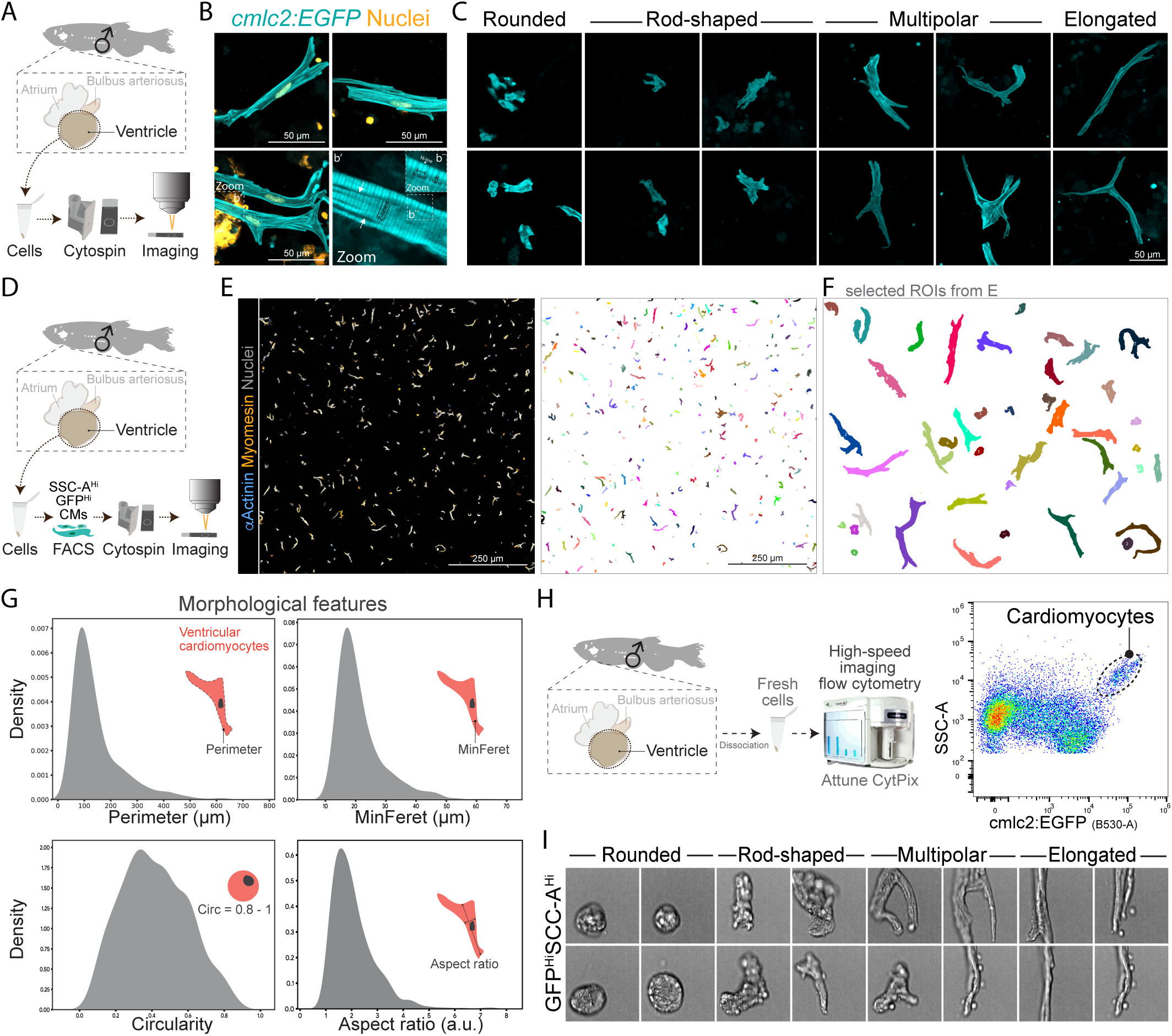
Substantial cell size and morphological variation of adult zebrafish ventricular cardiomyocytes. (A) Outline of adult male zebrafish ventricle isolation, dissociation, cytospin, and subsequent laser confocal imaging. (B) Representative laser confocal images of GFP+ cardiomyocytes showing intact and high-quality cells with visible sarco-meres. GFP fluorescence is shown in cyan, while nuclei are shown in yellow. (C) Representative laser confocal images of GFP+ cardiomyocytes showing intact cells with variable morphological shapes and sizes. Four main morphological features are highlighted, including rounded, rod-shaped, multipolar, and elongated cardiomyocytes. (D) Outline of cell isolation and FACS of GFP+ ventricular cardiomyocytes from adult male zebrafish. (E) Representative laser confocal images of FACS-isolated cardiomyocytes showing α-Actinin and myomesin staining. α-Actinin staining is shown in cyan hot, myomesin in orange hot, and nuclear staining in grey. The image on the right shows a resulting mask from the staining image on the left using Fiji. (F) Masks of randomly selected cardiomyocytes from (E) showing marked differences in the morphology and shape of cardiomyocytes. (G) Quantitative comparisons of perimeter, MinFeret, Circularity, and Aspect Ratio descriptors are shown (each group representing N = 6), revealing heterogeneity in male cardiomyocyte size and shape. (H) Outline of adult male zebrafish ventricle isolation, dissociation, and live-cell acoustic flow cytometry and bright field imaging of the GFP^Hi^SSC-A^Hi^ cardiomyocyte cluster. Cells were run without fixation. (I) Representative bright field images of GFP^Hi^SSC-A^Hi^ cardiomyocytes obtained using Attune CytPix acoustic focus.

### Morphometrics of Adult Zebrafish Ventricular Cardiomyocytes

To move beyond qualitative assessment and further characterize the size and morphological heterogeneity of cytospun ventricular cardiomyocytes, we quantitatively analyzed several shape descriptors—perimeter, minimum Feret diameter (MinFeret), circularity, and aspect ratio—from merged α-Actinin/Myomesin immunofluorescence images (**Fig. 4G, Fig. S6A**). High-throughput analysis of cell perimeter, representing total cell boundary length, revealed a unimodal but positively skewed distribution (**Fig. 4G, Fig. S6B, C**). This indicates that while most cardiomyocytes had smaller perimeters, a subpopulation displayed much larger perimeters, resulting in a long rightward tail. Median cardiomyocyte perimeter was approximately 100 µm.

Similarly, the MinFeret parameter, which reflects minimum cell width, showed a comparable right-skewed distribution with a median of 19 µm (**Fig. 4G**). In contrast, circularity—an index of cell roundness—followed a near-Gaussian distribution with mean and median values around 0.4, indicating a broad and asymmetric range of cell shapes (**Fig. 4G, Fig. S6C**). Aspect ratio, a measure of cell elongation, also exhibited a skewed distribution, further emphasizing the diversity in cardiomyocyte size and morphology (**Fig. 4G**).

To rule out the possibility that this morphological variability was an artifact of sample processing (e.g., fixation, cytospin, or shear stress during FACS), we employed acoustic-assisted flow cytometry with live-cell high-speed imaging of GFP^Hi^/SSC-A^Hi^ ventricular cardiomyocytes using the *AttuneCytPix* (**Fig. 4H**). Real-time imaging of unfixed, live dissociated ventricular cardiomyocytes from adult male *cmlc2:EGFP* zebrafish confirmed the presence of rounded and rod-shaped cardiomyocytes independent of fixation or cytospin procedures (**Fig. 4I**). Imaging-assisted flow cytometry also validated the presence of large, multipolar, and elongated cardiomyocytes in live preparations (**Fig. 4I**).

Collectively, these findings underscore the substantial morphological and size heterogeneity of adult zebrafish ventricular cardiomyocytes. This comprehensive, high-throughput characterization not only confirms the structural integrity of isolated cells but also offers new insights into the architectural and cellular complexity of the zebrafish heart, supporting the utility of our isolation protocol for downstream morphological, structural, molecular, and functional analyses.

### Sustained Exercise Promotes Ventricular Cardiomyocyte Hypertrophy and Hyperplasia

Zebrafish are valuable models for studying exercise-induced muscle and cardiac adaptations^29–33,56^. While cardiac structure and function can be readily evaluated at the whole-organ level, the detailed functional morphology of individual cardiomyocytes remains uncharacterized. To address this, we subjected adult zebrafish to a validated endurance swim regimen (**Fig. 5A**). Specifically, 7-month-old adult males underwent 4 weeks of sustained swimming at 70% of group critical speed (6 hours/day, 5 days/week; n=8 per group; see **Materials and Methods**). Following exercise training, we first assessed cardiac remodeling using high-frequency echocardiography. Resting heart rate decreased by 20 bpm (130.2 vs. 110.2 bpm; p = 0.0245) (**Fig. S7A**), confirming that extended swimming recapitulates the hallmarks of physiological cardiac hypertrophy in adult zebrafish and validates our model for subsequent single-cell analyses.

**Fig. 5.**
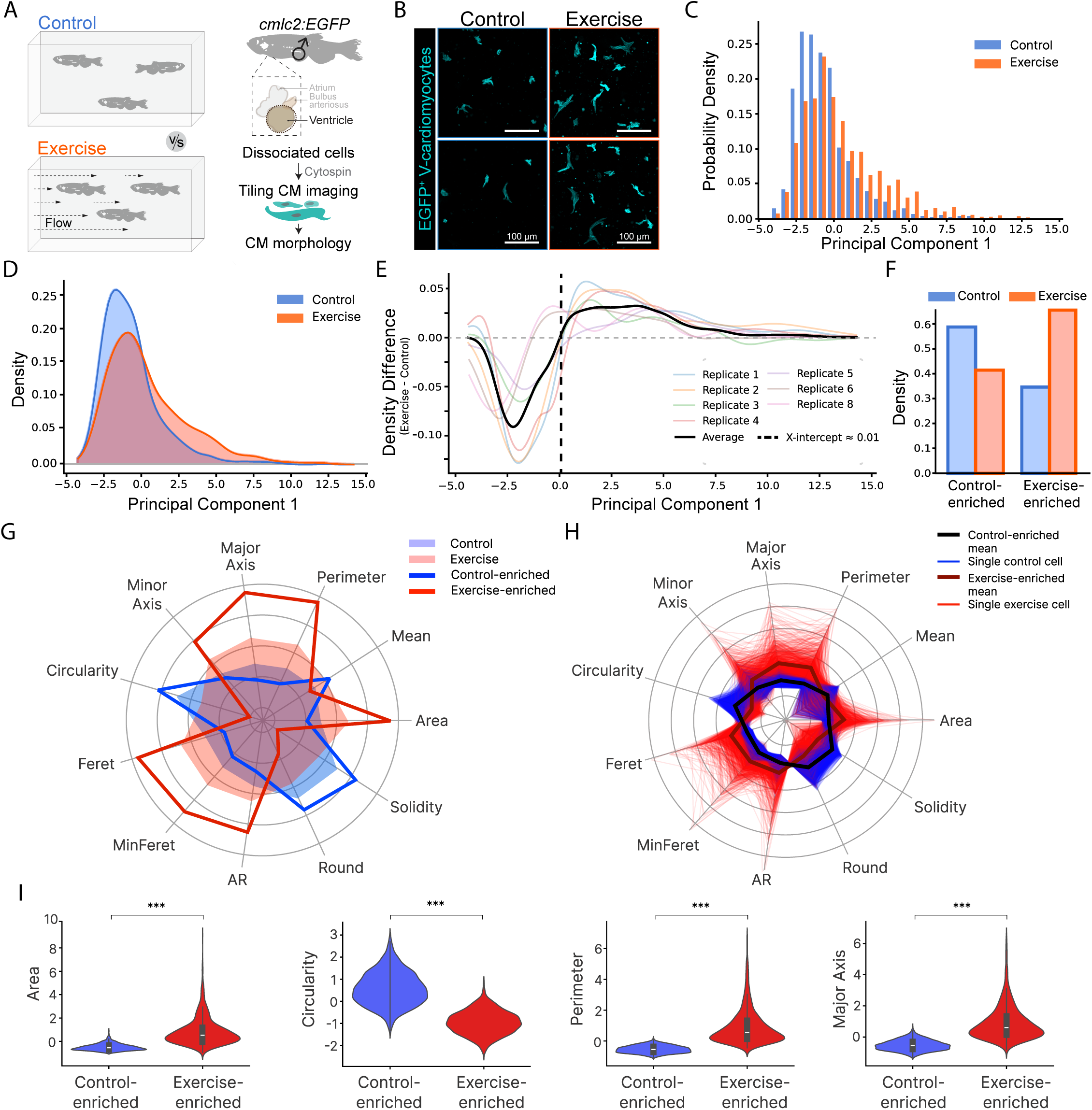
Swimming Exercise Induces Significant Morphological Remodelling of Cardiomyocytes. (A) Schematic of the experimental workflow. Cardiomyocytes (CMs) from control and exercise-conditioned cmlc2:EGFP transgenic fish were analyzed for morphology via imaging and flow cytometry. (B) Representative images of EGFP-labeled ventricular cardiomyocytes (V-CMs, cyan) from control and exercised fish. Scale bars: 100 µm. (C) Histogram of a 1-dimensional principal component analysis (PCA) using 11 normalised cell morphology features defines a comprehensive axis of phenotypic variation. Relative distributions of cells from Control and Exercise conditions are shown in blue and red, respectively. Dataset includes 2902 cells from 7 biological replicates. (D) Kernel density estimate (KDE) of Control and Exercise cell probability distributions relative to the PCA-based phenotypic axis emphasise differential enrichment of morphological phenotypes. (E) Difference curves between Exercise and Control KDEs (Exercise – Control). Mean of all data (solid black curve) and per technical replicate data (coloured curves). Vertical black dashed line marks the x-intercept, where relative enrichment shifts from control to exercise. Enriched cell populations (subsets) defined using this value. Control vs exercise enrichment population defined above and below this value respectively. (F) Column graph depicting enrichment of Control and Exercise condition cells within each defined subset. (G) Radar plot displaying mean phenotypic profiles for CMs prior to and post-subsetting, highlighting differences between control-enriched and exercise-enriched subsets at the population level. The plot includes 11 features derived from PCA. Feature values were z-score normalised. AR: aspect ratio. (H) Radar plot displaying single-cell phenotypic profiles from control-enriched and exercise-enriched subsets. Feature values were z-score normalised. (I) Violin Plots visualising the feature distribution of values for control- and exercise-enriched subsets. Features exhibit high variance between subsets, as determined in radar plot analyses. Annotated p-values were calculated using t-test.

Using the *cmlc2:EGFP* line for visualization (**Figure 5A**), we analyzed adult GFP⁺ dissociated ventricular cardiomyocytes (and/or co-stained with cTnT-A647/PE and/or α-Actinin-Vio^®^ R667) by tiling confocal microscopy and semi-automated image analysis. This approach revealed a 2.5-fold increase in cardiomyocyte yield following exercise (**Fig. S7B, C**). Qualitative examination indicated that individual cardiomyocytes from exercised fish appeared larger than those from controls (**Fig. 2B, Fig. S7B, C**). To quantify this morphological change, we performed one-dimensional principal component analysis (PCA) on 11 normalized cell features (Major and Minor axis, Circularity, Feret, MinFeret, Aspect Ratio [AR], Roundness, Solidity, Area, Mean, and Perimeter) from a dataset comprising 2,902 cardiomyocytes across 7 biological replicates. The PCA yielded a comprehensive axis of phenotypic variation that separated cardiomyocytes from control and exercised groups (**Fig. 2C**). Kernel density estimation (KDE) further emphasized differential enrichment of morphological phenotypes between groups (**Fig. 2D**), indicating that exercised hearts exhibit larger cardiomyocytes. These distinct shifts in cellular distributions prompted definition of phenotypically enriched cell subsets: difference curves (Exercise – Control) revealed a transition point along the PCA axis (**Fig. 2E**)—the x-intercept—where relative enrichment shifted. This defined "control-enriched" and "exercise-enriched" cell populations. Analysis confirmed a clear enrichment of control group cardiomyocytes within the control-defined subset, and, conversely, enrichment of exercise group cardiomyocytes within the exercise-defined subset (**Fig. 2F**).

To further characterize these populations, we visualized z-score normalized mean phenotypic profiles as radar plots. Across all PCA-derived features, exercised cardiomyocytes were significantly enlarged, thickened, and elongated compared to controls (**Fig. 2G**), with differences most evident between the exercise- and control-enriched subsets. At the single-cell level, individual radar plots reinforced these distinctions, revealing marked heterogeneity within each group while maintaining a clear phenotypic divergence (**Fig. 2H**). To highlight the most striking morphological differences, we generated violin plots of high-variance features (Area, Circularity, Perimeter, Major Axis) in control-versus exercise-enriched subsets (**Fig. 2I**). Unpaired t-tests confirmed highly significant increases in each [p<0.001]. Blinded quantification showed exercise induced an ∼15% increase in cell perimeter, comparable rises in MinFeret diameter and AR, and a ∼20% reduction in circularity—reflecting a shift toward more elongated shapes (**Fig. S7D**). Together, these results demonstrate that prolonged swim training induces robust, quantifiable eccentric hypertrophic remodeling of adult zebrafish cardiomyocytes, highlighting the sensitivity of our dissociation-based imaging and analysis pipeline.

Given the yield and quality of recovered cardiomyocytes, we next assessed cross-modality of the imaging pipeline in combination with flow cytometry. Using *cmlc2:EGFP;cmlc2:DsRed2-nuc* double-transgenic zebrafish, we found that GFP⁺DsRed2⁺ double-positive cardiomyocytes exhibited significantly higher forward scatter area (FSC-A, an indicator of cell size and complexity) and were recovered in 2.5–3 times greater numbers (**Fig. S7E–G**). This suggests that sustained exercise promotes both cardiomyocyte hypertrophy and hyperplasia. Overall, our approach—combining cell isolation with advanced morphological analysis—provides robust, quantitative methods for investigating the physiological adaptation of adult zebrafish cardiomyocytes to exercise.

### Establishment of a FACS Pipeline to Profile Cardiac Cells Using *cmlc2:EGFP;flk1:DsRed2* Double-Transgenic Zebrafish

Despite the widespread use of zebrafish as a model for cardiac biology, relatively few studies have comprehensively profiled adult cardiomyocytes (CMs) or endothelial cells (ECs) using bulk, in-depth molecular approaches. Notably, Kikuchi et al. (2011)^10^ assessed mRNA levels of retinoic acid signaling components using RT-qPCR from 15,000–20,000 DsRed2⁺GFP⁻ ECs or DsRed2⁻GFP⁺ CMs. Building on this, we aimed to determine whether global transcriptomic and proteomic landscapes could be profiled from much smaller isolated populations of cardiomyocytes and their more abundant endothelial neighbors. A key goal was to establish a workflow compatible with the inherently limited number of cardiomyocytes that can be practically isolated from about two ventricles.

To enable cell-type-specific proteo-transcriptomics, we first established a FACS protocol to co-purify CMs and ECs from the ventricles of adult *cmlc2:EGFP;flk1:DsRed2* double-transgenic zebrafish (**Fig. 3A, Fig. S8A**). Flow cytometric analysis of dissociated live ventricular cells clearly segregated the two target populations: flk1:DsRed2⁺ ECs (Cluster 1) and cmlc2:EGFP⁺ CMs (Cluster 2) (**Fig. S8B**). Post-sort imaging confirmed the identity of sorted flk1:DsRed2⁺ ECs as non-cardiomyocytes (**Fig. S8C**). Intriguingly, the cmlc2:EGFP⁺ CM population exhibited a broad range of DsRed2-like fluorescence in the yellow-green channel.

**Fig. 6.**
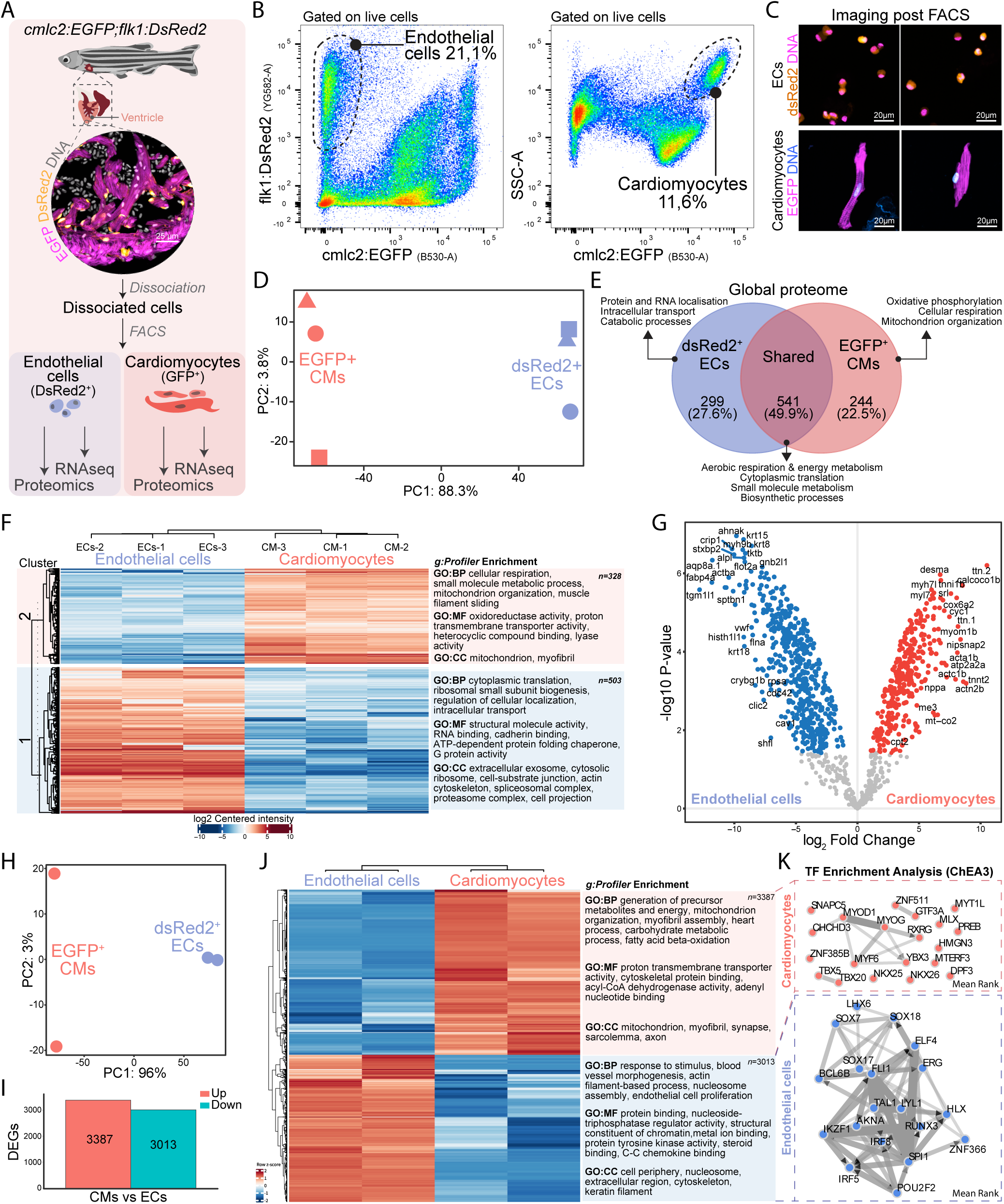
Low-input quantitative proteome and transcriptome profiling of adult zebrafish cardiac cells. (A) A shematic showing the strategy to perform dual-population FACS followed by proteomics and total RNA-seq using double transgenic reporter zebrafish. EGFP and dsRed2 channels were imaged using z stack confocal microscopy. DNA staining was also visualized. (B) Flow cytometry of dissociated cmlc2:EGFP; flk1:DsRed2 ventricles showing gating strategy. Endocardial cells were sorted by DsRed2 fluorescence; cardiomyocytes by GFP^Hi^SSC-A^Hi^. (C) Confocal images of FACS-purified populations confirm separation: DsRed2^+^ endocardial cells are smaller than GFP^+^ cardiomyocytes. (D) MaxQuant analysis identified 1084 reproducibly quantified proteins across the two cell types (ECs and CMs). (E) Proteome PCA shows distinct and significant clustering of endothelial cells (ECs) and cardiomyocytes (CMs) mainly along the PC1 axis. (F) Heatmap of 831 significantly differentially expressed proteins with GO enrichment (BP, CC, MF) indicating functional divergence. (G) Volcano plot of protein expression: red = upregulated in CMs, blue = upregulated in ECs (Log2FC > 1, -Log10 p < 0.05). (H) Transcriptome PCA shows clear separation of ECs and CMs along the PC1 axis. (I) Bar graph of differentially expressed genes: up/down in CMs vs. ECs. (J) Transcript heatmap with GO enrichment highlights functional differences. (K) Network of enriched transcription factors suggests key regulators of cell-type-specific expression, which are different between CMs and ECs.

To investigate this further, we sub-gated the cmlc2:EGFP⁺ population into DsRed2^Negative^, DsRed2^Low/Mid^, and DsRed2^High^ fractions and characterized them with immunofluorescence for the CM marker, TNNT2/cTnT. Cytospin preparations and confocal imaging confirmed that all three fractions contained bona fide cTnT-positive CMs (**Fig. S8D**). Critically, our FACS gating strategy, together with confocal microscopy, corroborated the lack of cellular doublets or aggregates, suggesting that the observed DsRed2 signal was contained *within* or *around* individual cardiomyocytes. In several cases, DsRed2 signal appeared as distinct intracellular puncta, with the majority of cardiomyocytes (∼65%) being DsRed-Negative, while smaller subsets displayed low/mid (∼25%) or high (∼5-10%) dsRed signal (**Fig. S8E**). These fluorescent puncta could alternatively represent extracellular vesicles (EVs) that retain RFP from their parent cells. Such vesicle-mediated transfer between ECs and CMs has been previously reported in the adult zebrafish heart^57^.

### FACS-Based Low-Input Analysis of Ventricular Cardiomyocyte and Cardiac Endothelial Cell Proteomes

Having established a robust FACS-based workflow for isolating CMs and ECs from *cmlc2:EGFP;flk1:DsRed2* zebrafish, we next sought to define the proteo-transcriptomic landscape of adult ventricular CMs and ECs. We performed untargeted LC-MS/MS proteomics and total RNA sequencing on FACS-purified populations from adult male hearts (n=3 biological replicates for proteomics; n=2 for transcriptomics) (**Fig. 3A–C, Fig. S9A**). To address the limited cell numbers obtainable per ventricle, particularly for CMs, we employed pooling strategies, typically combining 2–4 ventricles per biological replicate (see **Materials and Methods**).

Our low-input LC-MS/MS approach enabled global proteomic characterization from only 12,500 cells per sample, robustly quantifying 600–750 proteins per replicate with high quantitative consistency and no detected outliers (**Fig. S9B**). The close concordance in protein identifications underscores the technical reliability of our workflow and establishes a strong foundation for assessing differential protein expression.

Quality control confirmed both the integrity of the data and the distinct identity of the cell types. Pearson correlation and hierarchical clustering clearly separated samples into two primary branches corresponding to CMs and ECs, with very high intra-group correlation (r ≈ 1) and low inter-group correlation (r ≈ 0) (**Fig. S9C**). PCA further supported this molecular separation: the primary axis of variation (PC1) accounted for 88.3% of differences between cell types, while PC2 explained 3.8% (**Fig. 3D**). Thus, biological differences, rather than technical variability, accounted for the principal source of variation.

Following protein imputation (see **Materials and Methods**; **Fig. S9D**), label-free quantification (LFQ) analysis identified 840 proteins in ECs and 785 in CMs (**Fig. 3E**). Substantial proteome overlap was observed, with 541 proteins (∼50%) common to both cell types and enriched in core biological processes such as energy metabolism and protein translation. Additionally, 299 (27.6%) and 244 (22.5%) proteins were unique to ECs and CMs, respectively (**Fig. 3E**). Differential expression analysis (log₂FC > 1, adjusted P-value < 0.05) identified 503 proteins upregulated in ECs and 328 in CMs (**Fig. 3F, G**). Functional analysis of these protein sets revealed highly specialized roles for each cardiac cell type.

### Cardiomyocyte Enriched Proteome: An Electromechanical Powerhouse

The 328 proteins upregulated in cardiomyocytes define a striated cell highly specialized for powerful, rhythmic electromechanical activity. GO enrichment analysis using *g:Profiler*^58^ and *STRING*-based^59^ protein-protein interaction (PPI) networks revealed two major functional modules supporting this specialization: an advanced contractile apparatus and the high-capacity metabolic engine required to fuel it (**Fig. 3F, G, Fig. S9E**).

As expected, the cardiomyocyte proteome was dominated by proteins comprising the sarcomeric and contractile machinery. This included the giant structural proteins titin (Ttn.1 and Ttn.2), Z-disc anchor alpha-actinin-2 (Actn2)^54^, ldb3a/ZASP, and Z-disc linking proteins (Myoz2a, Myoz2b), and M-band components like myomesin (Myom1b, Myom2a). The structural framework was further reinforced by intermediate filament desmin (Desma), alongside core myofilament proteins including cardiac actin (Actc1b), myosin light and heavy chains (Myl7, Myh7), the troponin complex (Tnnt2, Tnnc1a), and tropomyosin (Tpm4a, Tpma) (**Fig. 3F, G, Fig. S9E, F**).

This contractile apparatus is supported by a robust enrichment of proteins driving high-energy metabolism. We observed upregulation throughout the metabolic pipeline: from cytoplasmic malate dehydrogenase (Mdh1), Me3, and enzymes for fatty acid β-oxidation (Cpt2, Cpt1ab, Acadm, Hadh)—the heart’s primary fuel source —to all principal components of the TCA cycle (Cs, Aco2, Idh2). This feeds into elevated mitochondrial oxidative phosphorylation (OXPHOS), as indicated by increased expression of subunits from all electron transport chain (ETC) complexes, including NADH dehydrogenase (Nduf family), cytochrome c oxidase (Cox family), enzymes for ubiquinone biosynthesis (Coq3, Coq5), and ATP synthase (Atp5f family) (**Fig. 3F, G, Fig. S9E, F**).

Supporting this metabolic output, proteins governing mitochondrial architecture were also upregulated. These included components of the MICOS complex (Immt, Micos10, Chchd3), which organize cristae junctions for efficient electron transport; Opa1 and Opa3, master regulators of mitochondrial fusion and cristae shape; and Tufm, which regulates mitochondrial protein synthesis. Notably, this metabolic engine is functionally coupled to the contractile apparatus via the creatine kinase system (Ckma), facilitating rapid transfer of high-energy phosphates from mitochondria to the sarcomere (**Fig. 3F, G, Fig. S9E, F**).

The cardiomyocyte proteome was also enriched for components essential to excitation-contraction coupling and calcium signaling, forming stable, interconnected PPI networks. These included the ryanodine receptor (Ryr2b) and calsequestrin-2 (Casq2), crucial for calcium handling, as well as proteins involved in mechanosensing, such as Muscle LIM Protein (Csrp3), the hormone precursor for atrial natriuretic peptide (Nppa) and the sarcolemma associated protein (Slmapa) (**Fig. 3F, G, Fig. S9E, F**).

### Endothelial Enriched Proteome: A Biosynthetic and Cell-Cell Interactive Hub

To more deeply characterize the molecular signature of cardiac ECs, we performed GO analysis on the 448 differentially expressed proteins (DEPs) significantly upregulated in this population, as well as PPI network visualization (**Fig. 3F, G, Fig. S9G, H**). These analyses revealed a proteome specialized for high biosynthetic activity, dynamic structural regulation, endocrinological and intercellular signaling, and the maintenance of a selective vascular barrier.

A significant portion of the upregulated proteome comprised core components for biosynthesis and protein quality control. This included a large number of ribosomal proteins (e.g., Rpl5b, Rps3a, Rplp0), elongation factors (Eif3s10, Eif4a1a, Eif4a3, Eif6), and molecular chaperones from the Hsp90, Hsp70, and CCT families (Hsp90ab1, Hsp90b1, Hspa5, Hspa8, Cct2, Cct5), underscoring the high capacity of ECs for protein synthesis and folding (**Fig. 3F, G, Fig. S9G, H**). Complementing these, numerous proteasome subunits (Psma5, Psmc2, Psme1) were also enriched, highlighting robust protein clearance and turnover mechanisms.

The cardiac EC proteome was rich in proteins dedicated to maintaining the dynamic cytoskeleton and cell adhesion. This included key regulators of the actin cytoskeleton (Cfl1, Arpc1a), components of the microtubule network (Tuba1b, Tubb2b), and proteins involved in specialized cell-cell and cell-matrix junctions. Notably, we identified core components of tight junctions (Tjp1a) and integrin-based focal adhesions (Itgb1b, Tln1/Talin-1, Ilk) (**Fig. 3F, G, Fig. S9G, H**).

Specialized endothelial structures and definitive lineage markers were strongly enriched, including caveolae components caveolin-1 (Cav1) and cavin family members (Cavin1b, Cavin2a), which are central to endothelial transcytosis and signaling^60,61^. The presence of von Willebrand factor (Vwf) unequivocally confirmed endothelial identity, while other endothelial-associated membrane proteins, such as Podxl/podocalyxin, were also enriched (**Fig. 3F, G, Fig. S9G, H**).

Metabolically, the EC protein profile highlighted an emphasis on glycolytic enzymes (Pgk1, Pkma) and proteins mediating redox homeostasis (Sod1, Prdx2, Txn). The complexity of endothelial gene regulation was reflected by the enriched RNA binding and splicing factors (Hnrnpl, Srsf1b, U2af2a). Finally, active cell signaling and trafficking were supported by the upregulation of small GTPases (Rac1b, Cdc42) and vesicular transport proteins from the Rab family (**Fig. 3F, G, Fig. S9G, H**).

In summary, while the endothelial dataset contains many ubiquitous housekeeping proteins, the specific enrichment of proteins involved in barrier formation, caveolae-mediated transport, and key markers such as Vwf provides a robust proteomic blueprint of the zebrafish cardiac ECs—one not previously available.

### FACS-Based Low-Input RNA-seq Reveals Distinct Transcriptional Landscapes in Cardiomyocytes and Endothelial Cells

Building on our global proteomics, we next applied our FACS workflow to transcriptomic profiling. A key challenge was extracting sufficient high-quality RNA from the limited cell numbers obtainable from zebrafish ventricles (**Fig. S10A**). Remarkably, as few as 7,000 DsRed2⁻GFP⁺ CMs yielded approximately 20 ng of high-quality RNA (RIN > 8.3) (**Fig. S10B**). While ECs are more abundant in uninjured hearts, they contain less RNA content per cell; thus, we FACS-purified 15,000 and 30,000 DsRed2⁺GFP⁻ cells, consistently recovering ∼27–35 ng of RNA with excellent integrity (RIN=9.3) (**Fig. S10B**). Paired-end total RNA-seq yielded at least 40 million raw counts per sample (**Fig. S10C**), and correlation matrix analysis and clustering clearly segregated samples by cell type: CMs and ECs (**Fig. S10C**). Metagene body normalization and *Picard*-based metrics confirmed even coverage with no 5’ or 3’ bias (**Fig. S10D**). Reads mapped to a wide away of RNA biotypes, including 22,873 coding genes, 1,476 lincRNAs, 321 pseudogenes, 114 miRNAs, 146 snoRNAs, 9 scaRNAs, and other RNA forms (**Fig. S10E**), underlining the complexity of our datasets. Normalized data distributions were consistent across samples (**Fig. S10F**).

PCA revealed that PC1 explained 96% of the variance between cell types, whereas PC2 explained only 3%, indicating that biological difference—not technical variability—was dominant, as previously observed in our proteomics data (**Fig. 3H**). Stringent DESeq2-based differential gene expression analysis (log₂FC > 2, adjusted p ≤ 0.05) identified 3,387 genes upregulated and 3,013 genes downregulated in CMs compared to ECs (**Fig. 3I**). A heatmap of all 6,400 differentially expressed genes (DEGs) confirmed the reproducibility and distinctness of these transcriptional states (**Fig. 3J**). These results demonstrate the robustness of our dissociation and FACS-based pipeline for generating high-quality RNA from limited cell numbers.

Functional annotation of CM-enriched DEGs revealed significant enrichment for Biological Process (BP) terms such as generation of precursor metabolites/energy, mitochondrial organization, myofibril assembly, heart process, and fatty acid elongation and β-oxidation. Molecular Function (MF) terms included proton transmembrane transporter activity, cytoskeletal protein binding, acyl-CoA dehydrogenase activity, and adenyl nucleotide binding. Cellular Component (CC) terms encompassed mitochondrion, myofibril, synapse, sarcolemma, and axon (**Fig. 3J**). These profiles recapitulate expected cardiac functions—sarcomere structure, contraction, mitochondrial energy metabolism, and more.

Consistent with these functions, the most abundant transcripts in CMs included canonical sarcomeric genes (*ttn.2* and *ttn.1*/ENSDARG00000000563, *myh7*, *myh7l*, *myl9b*, *myom1*, *myoz2b*, *myot*, *mybpc3*, *tnni4a*, *nrap*, *lmod3*, *obscnb*/ENSDARG00000022101, and *synpo2lb*), striated muscle genes (*xirp1*, *desma*, and *spega*), excitation-contraction coupling components (L-type calcium channel *cacna1*, *ryr2*, *atp2a2a* (Serca2a), *pln1*/phospholamban, *jph1a*/junctophilin-1a, *srl*/sarcalumenin, *slc8a1a*/ncx1), cardiac ion and calcium influx (*fxyd3*, *Vdac2*, *Vdac3*), mitochondrial energetics genes (*atp5fa1*, *mt-co1*, *mt-co2*, *mt-co3*, *mt-cyb*, *mt-atp6*, *mt-nd1*, *mt-nd4l*, *ndufa4b*, *cox4i1*, *cox6b2*, *ckma*, *mdh1ab*, *acadvl*, and *aifm1*), and cardiac developmental regulators (*smarcd3b*, *nkx2-5*, *tbx5a*, and *tbx20*) underscore the maintenance of CM identity and function^62^. Additional highly expressed genes included *gapdh*, *cdh2*, *popdc2*, *eno3*, *vegfab*, *fat1a*, *dennd4a*, *mylk4b*, *hsd11b2*, *fhl1a*, *corin*, *atf3*, *bag3*, *slc25a5*, *scn5lab*, *kcnh6a* (hERG channel), *hcn2*, *hcn4*, *neurl2*, *vldlr*, *nav2*, and *mlip* (ENSDARG00000089920) (**Fig. S10F**, **table S1**), supporting the diverse functional roles of CMs. Consistent with prior reports that Hspb7 is indispensable for mammalian heart development and actin filament regulation^63^, our transcriptomics and proteomics data reveal that *hspb7* is also predominantly expressed in cardiomyocytes.

EC-enriched GO:BP terms included response to stimulus, blood vessel morphogenesis, actin filament-based processes, nucleosome assembly, and endothelial proliferation. GO:MF categories highlighted protein binding, nucleoside-triphosphatase regulator activity, structural constituent of chromatin, metal ion binding, protein tyrosine kinase activity, steroid binding, and C-C chemokine binding. GO:CC terms included cell periphery, nucleosome, extracellular region, cytoskeleton, and keratin filaments (**Fig. 3J**). The most abundant endothelial transcripts included master regulators of endothelial identity and vascular development such as SRY-box transcription factors (*sox5*, *sox7*, *sox11a*, *sox18*), *erg*, *fli1*, *bcl6b*, Kruppel-like factors (*klf2a*, *klf2b*, *klf4*, *klf7b*, *klf17*), and the VEGF receptors *kdr* and *kdrl* (**Fig. S10F**, **table S1**). Genes involved in vascular morphogenesis and angiogenesis were prominently represented, including *egfl7*, *lama4*, *rasip1*, and the extracellular matrix and basement membrane components *col1a1a*, *col6a2*, *dcn*, *hspg2*, *spock3*, *itga1*, and *col4a1*. The mechanosensory ion channels *piezo1*, *piezo2*, and *trpv4* were highly expressed, consistent with endothelial mechanosensing and mechanotransduction in response to blood flow changes (**Fig. S10F**, **table S1**)^64–66^.

Additionally highly expressed EC genes were associated with regulation of vascular permeability, adhesion, and inflammation (*cdh5*, *clic2*, *sele*, *ncam3*, *cldn5b*, *ptgs2a/b*, *serpine1*, *fgl2a*, *f8*/ENSDARG00000101385), retinoic acid signaling (*aldh1a2*), guidance cues (*sema3ab*, *notch1b*, *notch1*), tetraspanin (*cd63*), actin binding (*flna*, *coro1ca*, *coro1cb*, *syne1*, *phactr4b*), cytoskeletal remodeling (*marcksl1a*, *dock6*, *iqsec1b*, *shroom4*), *krt8*, *krt18a.1*, *ahnak*, *epas1b*/Hif-2α, calcium signaling (*ramp2*), and small GTPase regulation (*agap1*, *kank4*, *dlc1*, *arhgap29a*/ENSDARG00000026329). Notably, *plk2b*, *yrk*, *slc6a4a*, *mmrn2a*, *stab1*, *aqp8a.1*, *cd74b*, *grb10a*, *akap12b*, *heg1*, *ahnak*, *errfi1a*, *flt1*, *ets2*/ENSDARG00000103980, and ENSDARG00000031658 (the human KIAA1217 ortholog) were also abundantly expressed. Importantly, many of these genes are, to our knowledge, described here for the first time in teleost endothelial cells, expanding the catalog of endothelial markers and functional candidates in the zebrafish heart.

To define regulatory networks underlying cell identity, we performed ChEA3-based^67^ transcription factor (TF) enrichment analysis on DEGs (**Fig. 3K**). In CMs, we identified core cardiac TFs including Myod1, Myog, Myf6, and Myt1l, which are known regulators of muscle differentiation and contractile function. Additionally, TFs such as Nkx2-5, Nkx2-6, Tbx5, and Tbx20, which are critical for cardiac development and chamber specification, were prominently enriched. Other notable cardiomyocyte-enriched TFs included Mterf3, Hmgn3, Znf511, Gtf3a, and the homeobox transcription factor Chchd3, reflecting the complex GRNs machinery required for maintaining cardiac contractile and metabolic programs (**Fig. 3K**). In ECs, we observed enrichment for the SOX family (Sox7, Sox17, and Sox18) and ETS family (Erg, Elf4, and Fli1)—master regulators of endothelial gene expression and vascular stability^68–71^. Additionally, Tal1, Lyl1, Runx3, Irf8, Spi1, and the zinc finger proteins Hlx, Znf366, Ikzf1 and Pou2f2 were identified (**Fig. 3K**), consistent with established roles in endothelial specification, proliferation, and vascular remodeling (**Fig. 3K**) Collectively, these profiles demonstrate that CMs and ECs are governed by distinct GRNs that reinforce their specialized functions within the vertebrate heart.

### Multi-Omics Integration and Identification of a Core Cardiomyocyte Proteome: Energy Production and Contraction

To define a CM and EC core proteo-transcriptomic map and determine key proteins driving CM and EC function and adaptation, we integrated DEPs from LFQ proteomics with bulk RNA-seq, using self-supervised machine learning (**Fig. 7A**, **Materials and Methods**). In cardiomyocytes, this analysis identified a high-confidence core set of 308 proteins/genes, each showing both abundance and active transcription. GO enrichment analysis and PPIs showed that the core proteome—networked strictly on physical interactions—is exquisitely optimized for two interdependent functions: aerobic respiration (energy production) and muscle contraction (force generation).

**Fig. 7.**
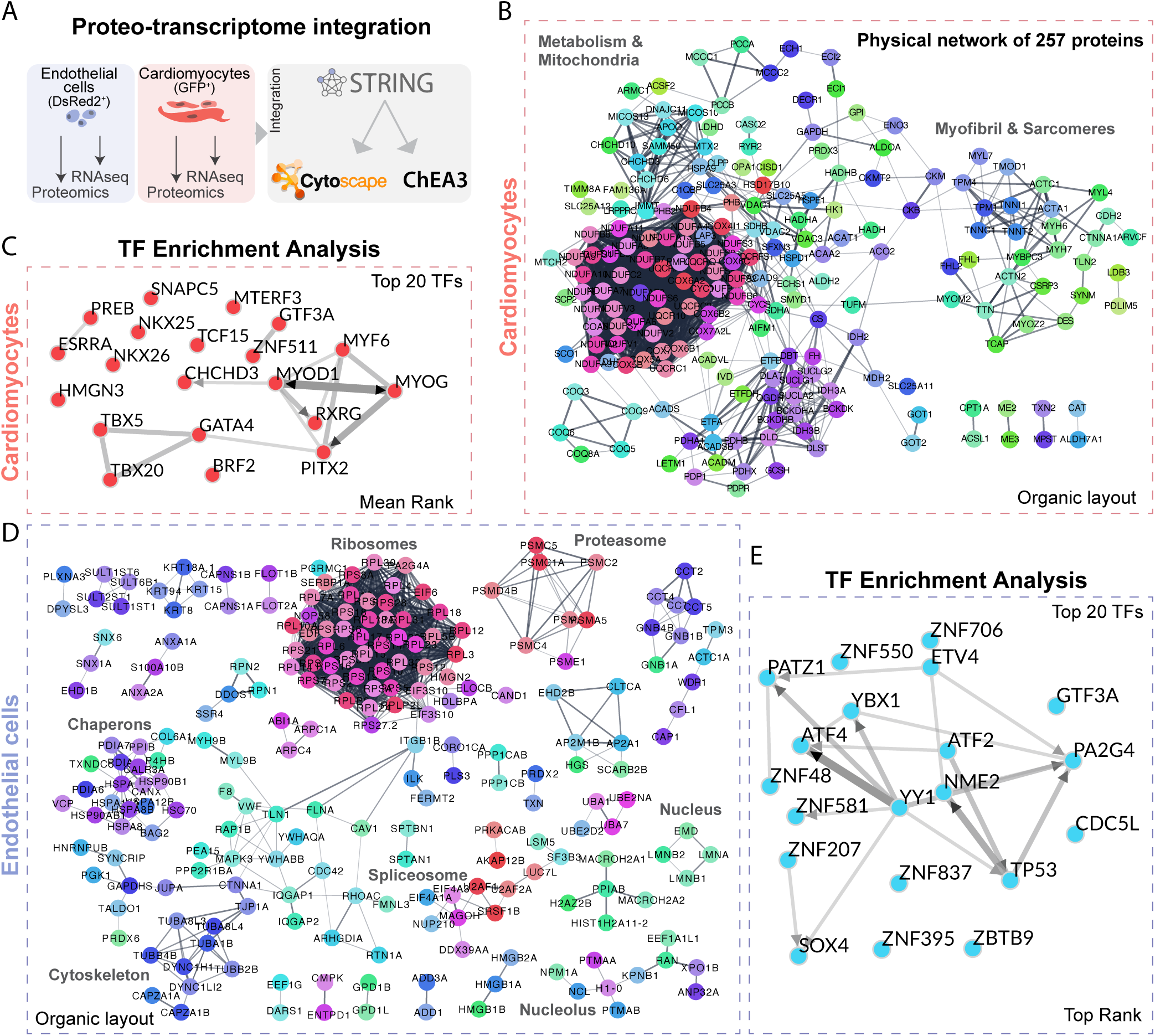
Functional integration of proteomic and transcriptomic data to map protein–protein interactions. Schematic of the dual-population workflow. cmlc2:EGFP^+^ cardiomyocytes and flk1:dsRed2^+^ endothelial cells are isolated by FACS, subjected to proteomic and transcriptomic profiling, and then integrated for multi-omic network analysis. (A) Cytoscape view of the cardiomyocyte protein–protein interaction (PPI) physical subnetwork. Nodes represent proteins with both transcript and protein evidence; edges are medium-confidence (0.4) physical interactions from STRING-db. An organic layout is used to emphasize functional clusters. Singletons were removed. Meaning of network edges: evidence (line color indicates the type of interaction evidence), confidence (line thickness indicates the strength of data support). (B) Subnetwork of the top 20 transcription factors enriched in cardiomyocytes, identified from the integrated proteome–transcriptome dataset. These TFs are putative master regulators of cardiomyocyte-specific gene expression. (C) Cytoscape view of the endothelial cell PPI network, generated as in (B) but using the flk1:dsRed2^+^ dataset. Singletons were removed. (D) Subnetwork of the top 20 transcription factors enriched in endothelial cells, highlighting candidate regulators of endothelial-specific proteome–transcriptome expression. Both cell types had a PPI enrichment p-value < 1.0e-16.

BP enrichment was dominated by mitochondrial energy production. Top categories were cellular respiration (P-value = 1.4 x 10⁻⁶⁵), ATP synthesis coupled to electron transport (95 proteins), represented by the rate-limiting enzyme for glycolysis hexokinase 1 (Hk1), lactate dehydrogenase D (Ldhd), pyruvate dehydrogenase complex component X (Pdhx), dihydrolipoamide S-acetyltransferase (Dlat), isocitrate dehydrogenase (NADP(+)) 2 (Idh2), succinate-CoA ligase GDP/ADP-forming subunit alpha (Suclg1), malate dehydrogenase 2 (Mdh2), several subunits of the supramolecular complexes of the respirasome including the ETC (e.g., Ndufa4, Uqcrfs1, Cox5a), and mitochondrial remodeling (Opa1, Micos10, Micos13, Samm50), inner membrane mitophagy supra-macromolecular receptors (Phb2/prohibitin 2) and Phb1/prohibitin 1, cytochrome C (Cyc1, Cycs), Mtx2/Metaxin 2, Vdac2, and mitochondrial pyruvate carrier 1 (Mpc1) (**Fig. 7B**).

Essential heart processes followed, including muscle contraction (P-value = 3.1 x 10⁻^41^, 48 proteins), notably core sarcomeric proteins (Ttn, Tnnt2, titin-cap/Tcap/telethonin, Actc1b, Myh7), as well as sarcoplasmic and membrane components (Ryr2, Slmap), and the calcium-binding protein Casq/calsequestrin 2 (**Fig. 7B**).

TCA cycle (P-value = 8.8 x 10⁻^18^) and fatty acid beta-oxidation (P-value = 2.5 x 10⁻^15^) pathways, both fueling OXPHOS, were also highly enriched, with key players including the pyruvate dehydrogenase complex (PDC) (Pdha1, Pdhb, Dlat, Dld), citrate synthase (Cs), isocitrate dehydrogenases (Idh2, Idh3a/b), alpha-ketoglutarate dehydrogenase (Odgh), the catalytic core of succinate dehydrogenase (Sdhb), mitochondrial matrix electron-transfer flavoproteins (Etfa, Etfb), Acadsb, and Carnitine O-acetyltransferase (Crat), and carnitine transferases (Cpt1a, Cpt2) (**Fig. 7B**).

CC analysis showed the core proteome is predominantly localized to the mitochondrion (P = 6.2 × 10⁻^98^, 188 proteins; 115 on the inner membrane—the ETC site). The second most enriched localization was the sarcomere (P = 1.1 × 10⁻^40^) and its sub-structures, such as the myofibril and Z-disc (**Fig. 7B**), highlighting essential contractile proteins (Mybpc3, Actn2b, Ttn.1).

MF enrichment emphasized oxidoreductase activity (P = 2.1 × 10⁻^45^), central to redox reactions in respiration (e.g., Ndufs1 and Sdhb), as well as ATP binding (P = 3.5 × 10⁻^32^, 78 proteins) and muscle structure maintenance (P = 1.8 × 10⁻^30^). Many actin-binding proteins (Actn2b, Tpm4a) further underscored the strong link to contractile sarcomeric function.

ChEA3-driven TF analysis of the 308-protein core revealed a network of 20 tightly interconnected TFs orchestrating cardiac gene expression (**Fig. 7C**). Master cardiac regulators (Nkx2-5, Nkx2-6, Tbx5, Tbx20, Gata4) myogenic factors (Myod1, Myog, Myf6), and additional TFs (Rxrg, Prf2, Pitx2, Chchd3, Znf511, Gtf3a, Mterf3, Preb, Snapc5, Tcf15, Hmgn3, Esrra) formed a rich, interconnected hub. All 20 TFs were strongly significant by mean rank, supporting coordinated control of the contractile and metabolic phenotype. Notably, metabolic regulators such as Esrra (a key factor in mitochondrial biogenesis and metabolism^72,73^ and Mterf3^74^ (mitochondrial transcription termination factor) emphasize the coupling of energy metabolism and contractile gene regulation. This network analysis illustrates how cardiomyocyte specialization arises from the coordinated action of cardiac developmental and metabolic TFs, paralleled by clustering of metabolic and myofibrillar/sarcomeric proteins (**Fig. 7B**).

### Multi-omics Integration and Identification of a Core Endothelial Proteome: Protein Synthesis and Cellular Structure

Overlapping abundant proteins with upregulated genes yielded a high-confidence set of 376 common proteins/genes. GO enrichment and PPI networks showed this proteo-transcriptomic map of zebrafish cardiac ECs is specialized for translation, cytoskeletal organization, protein folding/proteostasis, and glycolysis (**Fig. 7D**).

Translation was most significantly enriched (P-value = 8.5 x 10^-72^, 68 proteins), with abundant ribosomal proteins (Rpl5b, Rps3a, Rpl30, Rpl10a) and initiation factors (Eif3s10, Eif4a1a). Other significant terms included cytoskeleton organization (P-value = 1.4 x 10^-26^); Flna/filamin A and Vim/vimentin), protein folding (P-value = 3.0 x 10^-14^; Bag2, Hspa5/Bip, P4hb, Pdia3, Hsp90ab1, Hspa12a, Hspa8b, VCP/p97), vesicle-mediated transport, and glycolysis (**Fig. 7D**).

The ribosome was overwhelmingly overrepresented (P-value = 1.1 x 10^-81^, Fold Enrichment = 38.6). Many proteins also localized to the cytosol (P-value = 2.5 x 10^-63^), focal adhesions (P-value = 9.0 x 10^-29^) and plasma membrane, highlighting the importance of cell-matrix interactions and structural integrity in ECs. Other small represented but perhaps relevant proteins are associated with the nuclear structural maintenance and genome regulation including the nuclear lamina (Lmna, Lmnb1, Lmnb2), Histones (H1-0, H2az2b, Macroh2a1, Macroh2a2) and nucleolar proteins (Ncl, Npm1a) (**Fig. 7D**), linking these with high ribosomal production and content.

The MF analysis confirmed the findings from the other domains. Structural constituent of ribosome (P-value = 7.3 x 10^-85^), and RNA binding (P-value = 3.1 x 10^-49^) were most enriched (heterogeneous nuclear ribonucleoproteins [e.g., Hnrnpl, Hnrnpa1a]), with ATP binding and actin binding also prominent, consistent with high biosynthetic and structural activity.

ChEA3 analysis on the 376-protein endothelial core highlighted a distinct network of 20 top TFs dominated by proteins that orchestrate endothelial gene expression programs (**Fig. 7E**), such as zinc finger transcription factors (Znf706, Znf550, Znf48, Znf581, Znf207, Znf837, Znf395) and transcriptional regulators involved in RNA processing and chromatin remodeling. Key endothelial regulators included Ybx1 (Y-box binding protein 1), a multifunctional protein involved in transcription, translation, and mRNA packaging; Atf4 and Atf2 (activating transcription factors), regulating stress responses and cellular adaptation; and Nme2 (nucleoside diphosphate kinase), playing roles in transcriptional regulation and cellular differentiation (**Fig. 7E**). The presence of Sox4, a member of the SOX (SRY-related HMG-box) TF family known for roles in developmental processes and endothelial specification, further confirms endothelial identity. Additional enriched TFs included Tp53 (tumor protein p53), a master regulator of cell cycle and stress responses; Patz1 (POZ/BTB and AT hook containing zinc finger 1); Etv4 (ETS variant 4); Gtf3a (general transcription factor IIIA); Pa2g4 (proliferation-associated 2G4); Cdc5l (cell division cycle 5-like); Yy1 (Yin Yang 1); and Zbtb9 (zinc finger and BTB domain containing 9) (**Fig. 7E**). Together, these transcriptional networks align with the proliferative nature of coronary endothelial cells in adult zebrafish^43^.

The network connectivity analysis revealed that Znf581 and Ybx1 serve as central regulatory hubs, connecting multiple transcriptional pathways. Unlike the cardiomyocyte TF network, which was dominated by cardiac developmental regulators (Nkx2-5, Tbx5, Gata4)^62^, the endothelial TF network reflects the cell’s primary functions in biosynthesis, chromatin remodeling, and adaptive responses to environmental stimuli. The mean rank analysis demonstrated consistent enrichment across the dataset, with all top 20 TFs showing strong statistical significance.

This TF signature aligns with the functional specialization observed in the PPI networks and our proteo-transcriptome datasets (**Fig. 3**, **Fig. 7D, Fig. S9G, H**), showing distinct clustering of ribosomes (protein synthesis machinery), proteasomes (protein degradation), chaperones (protein quality control), cytoskeleton, spliceosome (RNA processing), and nuclear components. The enrichment of TFs involved in transcriptional regulation (Ybx1, Atf4, Atf2, Yy1), chromatin organization (multiple ZNF proteins), and stress responses (Tp53, Atf4) underscores the endothelial cell’s role as a metabolically active, biosynthetically intense, and environmentally responsive barrier. The prominence of zinc finger proteins suggests complex epigenetic regulation underlying endothelial plasticity and adaptation to hemodynamic forces.

## DISCUSSION

We developed and validated a robust, physiologically matched workflow for isolating, imaging, and molecular profiling of adult zebrafish and medaka cardiac cells. Our pipeline overcomes longstanding limitations in cell yield, viability, and molecular resolution, enabling precise FACS-based purification, comprehensive morphological characterization of cardiomyocytes, and multi-omics data from low-input ventricles. As a result, we generated the first integrated proteo-transcriptomic atlas of teleost heart cell types, providing new insights into their cellular, metabolic, and regulatory specialization.

Transitioning from mammalian-like dissociation at 33–37°C to the physiological range for teleosts (e.g., zebrafish at 26.5-28°C)^45^ proved pivotal, dramatically improving cell viability (>95%) and preserving morphology without increasing processing time. Although a prior adult zebrafish cardiac regeneration study used 28°C dissociation^9^, its analysis was restricted to immune cells, leaving cardiomyocyte quality and yield unknown. Elevated temperatures can induce heat-shock responses, alter gene expression, and compromise cellular architecture^75–77^; by maintaining physiological temperatures, we minimize stress-induced transcriptional noise and ensure our datasets reflect true biological signatures rather than methodological artifacts.

In *cmlc2:EGFP* hearts, the GFP^high^/SSC-A^high^ population represents viable, intact cardiomyocytes, whereas the GFP^high^/SSC-A^low^ population consists primarily of cellular debris— an often-missed distinction^39^. Failure to exclude debris can artificially inflate cell counts, introduce contamination, and compromise downstream analyses. We provide critical benchmarks for experimental planning, reporting yields of approximately 4,000–7,000 live cardiomyocytes per ventricle in young and mature adult male zebrafish, about half that number in aged males, and roughly double in adult male medaka. This age-dependent decline likely reflects reduced cardiomyocyte numbers and increased fibrosis. Our workflow offers a reproducible template across laboratories and underscores the value of microscopy-based validation coupled with flow cytometry to isolate high-quality teleost cardiomyocytes.

One of the most striking findings is the morphological diversity of adult zebrafish ventricular cardiomyocytes. While prior work typically describes them as "elongated"^55^ or "rod-shaped"^78,79^, our unbiased quantitative profiling revealed a spectrum, including rounded, rod-shaped, multipolar, and highly elongated forms. This diversity was observed in live, unfixed cells using acoustic-assisted imaging flow cytometry, ruling out processing artifacts. This heterogeneity may reflect functional specialization within the ventricle (trabecular^80^, primordial layer^81^, compact myocardium, or regional differences^78^), or differences in metabolic states, cell cycle, or maturation. The narrow, elongated morphology is conserved across most species^55^. Our morphometric framework establishes a foundation for linking physical phenotype to function and molecular signatures through single-cell omics.

Our analysis provides the first single-cell view of physiological cardiomyocyte hypertrophy in zebrafish. Four weeks of sustained swimming induced heart remodeling and decreased resting heart rate. Exercised cardiomyocytes exhibited ∼15% increases in perimeter and MinFeret diameter, ∼20% increase in length, and ∼20% decrease in circularity—all signatures of hypertrophic elongation. The increased cardiomyocyte yield suggests that both hypertrophy and hyperplasia contribute to exercise-induced growth^32^. Previous work showed swimming exercise stimulates cardiomyocyte proliferation without altering ventricular size/function^31^, and enhances proliferation after cryoinjury (∼4-fold)^31^, in contrast to mammals, where adult cardiomyocytes are largely post-mitotic and adaptation is primarily hypertrophic^82,83^.

The mechanisms driving this plasticity remain incompletely defined. Our platform integrates morphology with transcriptomic and proteomic profiling to identify the pathways, transcriptional programs, and metabolic shifts engaged by exercise. Clarifying why zebrafish cardiomyocytes retain proliferative potential, and how physiological stress augments it, may reveal targets for human cardiac repair. Future work should also test sex-specific responses, as current data largely derive from males or mixed-sex cohorts.

Our workflow resolves rare, dedifferentiated *gata4*⁺ cardiomyocytes after ventricular resection, capturing their expansion and morphological shift during regeneration. At 7dpi, *gata4*⁺ cardiomyocytes expanded ∼3–to 4-fold and assumed rounded morphology, consistent with dedifferentiation—a critical event in zebrafish heart regeneration^1,2,5^. This resolution enables FACS isolation of high-purity *gata4*⁺ populations for omics profiling to interrogate programs of dedifferentiation, proliferation, and redifferentiation. Comparative analyses between regenerative and non-regenerative species may identify key regulators of regenerative capacity with translational relevance^7,10,36,37,51,84^.

A central achievement is the integration of low-input proteomics and transcriptomics to assemble an atlas of adult teleost ventricular cells. From as few as 7,000–15,000 FACS-purified cells, we quantified 785 cardiomyocyte and 840 endothelial proteins and identified >6,000 DEGs separating the two populations. Strong within-group (r ≈ 1) and minimal between-group (r ≈ 0) correlations validate data quality and biological distinction. CMs were enriched for sarcomeric structure, contraction, and energy metabolism, whereas ECs were enriched for blood vessel morphogenesis, nucleosome assembly, mechanosensation, and vascular homeostasis.

Cd63, an established exosome/EV marker essential for vesicle biogenesis and cargo sorting, and intercellular communication^85^, shows pronounced enrichment in endothelial cells. This robust *cd63* expression indicates active EV production and secretion by the endothelium. Coupled with our observation of fluorescent dsRed2 puncta within cardiomyocytes, these findings support a model in which endothelial-derived vesicles mediate intercellular communication that underlies cardiac homeostasis and regeneration.

By integrating low-input proteomics and RNA-seq, we resolved the core molecular programs that distinguish CMs and ECs. In cardiomyocytes, mitochondrial proteins dominate the proteome—reflecting extraordinary energetic demands—with comprehensive representation of electron transport chain complexes, fatty acid β-oxidation enzymes, TCA cycle components, and ATP synthases that fuel rapid, rhythmic contraction.

The sarcomeric proteome was equally comprehensive, encompassing giant proteins like titin (Ttn.1, Ttn.2), Tcap/telethonin, Z-disc anchors (Actn2, ldb3a/ZASP, Myoz2), M-band components (Myom1b, Myom2a), Mybpc3, and all major contractile filament proteins (myosin heavy/light chains, actin, troponin, tropomyosin), highlighting key mechanotransduction nodes and structural assemblies. Metabolism and contraction are functionally coupled via the creatine kinase system (Ckma), facilitating rapid phosphate transfer from mitochondria to sarcomeres, and calcium-handling proteins (Ryr2, Casq2) for excitation-contraction coupling. TF enrichment analysis identified core cardiac developmental regulators (Nkx2-5, Tbx5, Tbx20) and myogenic factors (Myod1, Myog, Myf6) orchestrating these networks and maintaining cardiomyocyte identity. Together, these data define the cardiomyocyte as a highly specialized, metabolically intense contractile machine, providing a baseline for interpreting how CMs undergo rapid metabolic and structural switching in response to physiological challenges like exercise, injury or pathological stress.

The core endothelial signature included approximately seventy ribosomal proteins and abundant translation initiation factors, consistent with high protein synthesis supported by quality-control machinery (Hsp70/90 chaperones, proteasome components, disulfide isomerases). A reinforced cytoskeletal network (actin regulators Cfl1, Arpc1a; tubulins; vimentin; filamin) underpins structural integrity and hemodynamic responsiveness, while specialized membrane domains—caveolae (Cav1, Cavin1b, Cavin2a) for transcytosis and signaling^86^—and tight junctions complexes (Tjp1a/ZO-1, Shank3) sustain barrier function. Definitive endothelial markers (Vwf/von Willebrand factor and Podxl/podocalyxin) and mechanosensory ion channels (Piezo1, Piezo2, Trpv4) confirm identity and place mechanotransduction at the core of endothelial physiology. Notably, Piezo1 mediates endothelial inflammation^87^, making it a promising therapeutic target for atherosclerosis and cardiovascular disease. Our proteo-transcriptomic maps provide a cell-resolved resource to interrogate endothelial dysfunction and EC–CM crosstalk during homeostasis, regeneration, aging and drug- or exercise-induced remodeling. Transcription factor analysis revealed that this biosynthetic and barrier-forming phenotype is governed by SOXF group (Sox7, Sox17, Sox18)^88–90^ and ETS family (Erg, Fli1, Elf4)^68,70,71,91,92^, which are master regulators of vascular development and endothelial specification.

Beyond catalogs of individual genes and proteins, integrated protein-protein interaction (PPI) networks—prioritizing physical associations—uncover higher-order functional modules (**Fig. 7**). In CMs, networks centered on mitochondrial respiratory complexes, sarcomeric machinery, and calcium handling, while in ECs, networks centered on ribosomal and chaperone assemblies, proteasome complexes, nuclear/nucleolar, and focal adhesion components emerged. These architectures reveal how biosynthetic, structural, and signaling pathways are coordinated in a cell-type-specific manner, demonstrating that CMs and ECs are governed by fundamentally distinct GRNs reflecting their specialized physiological roles. Future integration of phosphoproteomics, lipidomics, metabolomics, and single-cell transcriptomics will further resolve the dynamic regulation of these molecular machines during development, regeneration, and disease.

By addressing long-standing technical barriers and delivering molecular atlases at cellular resolution, our platform establishes a new standard for teleost heart research. The extensive methodological detail, standardized gating strategies, cardiomyocyte morphometrics, and multi-omic datasets we provide will enhance reproducibility and facilitate cross-study comparison. While previous studies mapped bulk cardiac proteomes across species and chambers^93^, our cell-type-resolved approach—now essential in mammalian systems^94,95^—circumvents tissue heterogeneity and exposes previously hidden molecular programs. By generating datasets at cellular resolution, we illuminate the “*dark* protein-protein interactome”^96^, still poorly defined in fish. As single-cell and systems-scale approaches continue to advance, the foundation established here will drive discoveries spanning evolution, regeneration, and human cardiovascular health.

## LIMITATIONS OF THE STUDY

While our work represents a significant advance, several limitations and avenues for future improvement remain. First, our bulk RNA-seq and proteomics analyses average across cell populations, potentially masking rare cell types or subtle cell-state heterogeneity. Incorporating single-cell or single-nucleus RNA-seq/ATAC-seq would provide finer resolution and enable lineage or trajectory analyses, particularly during regeneration or disease. Second, our protocol currently pools 2–4 ventricles per replicate to obtain sufficient input; further optimization of ultra-low-input methods (e.g., Smart-seq3 or nanoPOTS proteomics), in combination with next-generation mass spectrometers, may eventually allow profiling at the level of individual hearts, thereby increasing throughput and reducing biological variability. Third, although we focused on ventricular cardiomyocytes and endothelial cells, other cardiac populations—such as fibroblasts, immune, and epicardial cells—remain underexplored. Extending our methods to these populations will be essential for achieving a complete cell-based molecular atlas of the teleost heart. Fourth, while our morphological profiling revealed substantial cardiomyocyte heterogeneity, functional studies—including patch-clamp electrophysiology, calcium imaging, or contractility assays on FACS-purified subpopulations—will be required to directly link morphology to function. Lastly, applying this workflow to models of cardiac injury, congenital heart disease, cardiomyopathies, or arrhythmia will be important for validating its translational utility and for uncovering mechanisms relevant to human cardiovascular medicine.

## MATERIALS AND METHODS

### Zebrafish and Medaka Husbandry and Strains

Wild-type and genetically modified zebrafish (*Danio rerio*) from Ekkwill (EK) and TE genetic backgrounds and medaka fish (*Oryzias latipes*) from Cab strain were maintained at the Victor Chang Cardiac Research Institute. Transgenic zebrafish lines used in this study were *Tg(myl7:EGFP)*^20^, *Tg(myl7:EGFP;flk1:DsRed2)*, Tg(-5.1cmlc2:DsRed2-nuc)^97^, and *Tg(kdrl:DsRed2)*^10^*. Myl7* was previously called *cmlc2*.Transgenic medaka used were *Tg(cmlc2:GFP*). Adult zebrafish were housed in 3.5 L tanks and adult medaka in 10 L tanks using the Z-Hub system (Aquatic Habitats), with a maximum of 5 fish per L. Fish were kept in recirculating, chlorine-free water at a temperature of 27 ± 1 °C, under a 14.5-hour light and 9.5-hour dark cycle, and were fed twice daily (pellets and artemia). All experimental procedures were conducted in accordance with protocols approved by the Garvan Institute of Medical Research/St Vincent’s Hospital Animal Ethics Committee (Animal Research Authority: AEC:22_25 and AEC:21_16) and the Australian National Health and Medical Research Council code of practice for the care and use of animals.

### Ventricular Tissue Dissociation and Single-Cell Isolation

Adult male zebrafish and medaka were used for ventricular cell isolation. Fish were euthanized and hearts harvested as per previous protocols^98^. Harvested hearts were cleaned of surrounding tissue and fat, and ventricular tissue was isolated from the rest of the heart and kept at 4°C in PBS1x during tissue collection. The ventricles were then opened halfway and transferred to 2 mL Eppendorf tubes maintained on ice. Pooled ventricles (two-to-four ventricles per tube) were washed with ice-cold FACS buffer (1× PBS, 2% heat-inactivated fetal bovine serum [FBS], 2 mM EDTA, pH 7.9) to remove residual blood. A one-step enzymatic dissociation protocol was employed^48^. Our custom-made *zebrafish dissociation buffer* (ZF3.0) was prepared by mixing equal volumes (1:1) of digestion buffer and papain buffer to a final volume of 50 mL. The digestion buffer (25 mL) was prepared in DMEM (Cat. No. 11995065, Thermo Fisher) containing L-glutamine, sodium pyruvate, and 4.5 g/L glucose, supplemented with collagenase II (265 U/mL; Cat. #LS004176, Worthington), bovine serum albumin (1%; Cat. #A7906, Sigma), dispase (0.2 U/mL; Cat. #07913, STEMCELL), and DNase I (0.050 mg/mL; Cat. #10104159001, Sigma-Aldrich). The papain buffer (25 mL) was prepared in the same DMEM formulation and contained papain (25 U/mL; Cat. No. 10108014001, Roche), N-acetyl-cysteine (2 mM), taurine (30 mM), and 2,3-butanedione monoxime (BDM; 10 mM). Taurine and BDM were added immediately before combining with the digestion buffer. Prior to use, the complete ZF3.0 dissociation buffer was diluted by mixing equal volumes (1:1) with 1% BSA. The ZF3.0 buffer was used within 6 months of preparation while stored at -20°C. Initially, 500 μL of ZF3.0 + 1% BSA (1:1 v/v) buffer was added per two ventricles. Tubes were incubated in a thermoregulated Eppendorf mixer (900 RPM) at 28°C for 30–45 minutes, with vigorous manual shaking (10-20 repetitions) every 15 minutes. Following complete dissociation, 1 mL of ice-cold FACS buffer was added to the dissociation tube, followed by gentle manual shaking. Tubes were centrifuged at 500 × g for 5 minutes at 4°C, and the resulting pellet was resuspended in 500-1,000 µL of cold FACS buffer.

### Live/Dead Staining and Sample Preparation

For viability assessment, an additional 500 µL of ice-cold FACS buffer was added to bring the total volume to approximately 1.0 mL. The suspension was centrifuged at 500 × g for 5 minutes at 4°C and the supernatant removed, leaving a final cell suspension of approximately 100 µL. Cells were stained with 500 µL of Live/Dead (L/D) mix containing Calcein-AM (0.5 µM), propidium iodide (PI, 1 µg/mL), DAPI (1 µg/mL) and Zombie Yellow dye (1:1000 dilution) for 10 minutes at 30°C. Calcein-AM was ommitted when using GFP transgenic fish. PI was omitted when using DsRed2/RFP transgenic fish. If cells were intended for FACS, 0.5 mL of FACS buffer was added post-staining. For cytospin preparations, 50–100 µL of the cell suspension was fixed with paraformaldehyde (PFA) to a final concentration of 2% (v/v), and cells were cytospun at 2,000 rpm using a cytocentrifuge. Flow cytometry samples were analyzed in a final volume of approximately 500-1,000 µL.

### Embryonic Tissue Dissociation and Single-Cell Isolation

*Tg*(*cmlc2:EGFP*) zebrafish embryos were maintained in E3 medium at 28.5 °C until visually staged. At 28.5 hpf, embryos were collected and pooled into 1.5 mL Eppendorf tubes, with approximately 20 embryos per tube. Embryos were manually dechorionated and washed with ice-cold FACS buffer to remove residual yolk and debris. For enzymatic dissociation, embryos were incubated in ZF3.0 buffer diluted 1:1 (v/v) with 1% BSA. Tubes were placed in a thermoregulated Eppendorf mixer (900 RPM) at 28°C for 30–45 minutes, with vigorous manual shaking (20 repetitions) every 15 minutes to facilitate complete tissue dissociation. Following dissociation, 1 mL of ice-cold FACS buffer was added to each tube, followed by gentle manual shaking. Samples were centrifuged at 500 × g for 5 minutes at 4°C, and the resulting pellet was resuspended in 500–1,000 µL of cold FACS buffer. For viability assessment, cells were stained with Live/Dead mix containing propidium iodide (PI, 1 µg/mL), DAPI (1 µg/mL), and Zombie Yellow dye (1:1000 dilution) for 10 minutes at 30°C. Flow cytometry analysis was performed using a CytoFlex flow cytometer. Calibration beads were used as size reference markers to aid in cell population gating. Data were analyzed using FlowJo Portal (version 10.8.1, BD). For embryonic samples, approximately 300,000–500,000 total events were captured per analysis.

### Zebrafish and Medaka Nuclei Isolation

Following ventricular dissection, single-cell ventricular suspensions were obtained as previously described from n=2 for zebrafish and medaka (3x ventricles per tube, totalling 2 tubes). The cell pellet was resuspended in 500 μL of lysis buffer (EZ Lysis Buffer, Nuclei EZ Prep, Sigma Aldrich, Cat# NUC101) supplemented with 0.2 U/μL Recombinant RNasin® Ribonuclease Inhibitor (Promega, Cat# N2511), then mechanically dissociated by pipetting up-and-down five times with a RAININ 200 μL BioClean Ultra UNV Pipette Tip (Mettler Toledo, Cat# 30389188) followed by five times with a 1000 μL tip. After incubation on ice for 3 minutes, nuclei were pelleted by centrifugation (500 g, 5 minutes, 4°C).

Nuclei were then resuspended in 1 ml resuspension buffer (1% Bovine Serum Albumin [BSA], Sigma Aldrich, Cat# A7906-50G in 1X DPBS [Gibco, Cat# 14190-144], supplemented with 0.2 U/μL RNasin® Ribonuclease Inhibitor), and centrifuged again under the same conditions. The final nuclear pellet was resuspended gently in ∼200 μL resuspension buffer and a 1:2500 dilution of VybrantTM DyeCycleTM Violet (V35003, 5 mM) and propidium iodide (1µg/mL) was added. The tubes were covered and taken for analysis by flow cytometry.

### Flow Cytometry Analyses and fluorescence-activated cell sorting (FACS)

Flow cytometry analyses were performed using a BD LSRFortessaTM X-20 Cell Analyzer and a BD FACSymphony A5 High-Parameter Cell Analyzer, both equipped with five excitation lasers (UV 355 nm, Violet 405 nm, Blue 488 nm, Yellow/Green 561 nm, and Red 633 nm). Freshly isolated cells were sorting using a BD FACSAria™ III Cell Sorter equipped with an ND2 filter and 100µm nozzle (specifically used for zebrafish large cardiomyocytes). Data were collected using BD FACSDiva™ Software. FSC-H versus FSC-A, FSC-H versus FSC-W, and SSC-H versus SSC-W cytograms were used to discriminate and gate out doublets/cell aggregates during sorting or analysis. Cells were first gated for singlets, followed by Live/Dead analysis based on CalceinAM (B530-A) fluorescence intensity, and propidium iodide (YG610-A) and DAPI (V450-A) exclusion. Our analyses revealed more than 95% of cells were single-cell preparations. We selected single cells that were CalceinAM positive and PI negative. Cells were sorted in 1% BSA (pH 7.4). For optimal DNA dye signal detection and Live/Dead analyses, an event concentration of <1,500 events/seconds was used, and 30,000-100,000 total events were recorded. For analysis of zebrafish embryos, around 300,000-500,000 events were captured. All flow cytometry data were analyzed using FlowJo Portal (version 10.8.1, BD) using macOS Monterey. Manual compensation was done only when exclusively required (e.g., DAPI vs GFP channel or GFP vs DsRed).

### Quantification of Cardiomyocyte Yields Across Adult Zebrafish Lifespan Animals and Age Groups

Male Tg(cmlc2:EGFP) zebrafish were grouped into three adult life stages: young adults (4-5 months post-fertilization, mpf), mature adults (6-10 mpf), and aged individuals (12-16 mpf). Fish were maintained as described above.

### Ventricular Dissociation and Single-Cell Preparation

Individual zebrafish were euthanized by tricaine overdose, and hearts were rapidly excised and transferred to ice-cold phosphate-buffered saline (PBS). Ventricles were carefully isolated from atria and bulbus arteriosus under a stereomicroscope. Single ventricles were dissociated using our optimized protocol as described above. Following dissociation, cell suspensions were resuspended in 1 mL of FACS buffer.

### Flow Cytometry Analysis and Cardiomyocyte Quantification

Flow cytometry was performed using both BD LSRFortessaTM X-20 Cell Analyzer and FACSAria™ III Cell Sorter. For each sample, the entire cell suspension volume was analyzed to enable absolute cell counting. Single-cell events were gated based on forward scatter area (FSC-A) versus side scatter area (SSC-A), followed by singlet discrimination using FSC-A versus FSC-height (FSC-H). Live cells were identified by exclusion of viability dyes as described above. GFP⁺ cardiomyocytes were identified as GFP^high^/SSC-A^high^ events to exclude cellular debris (GFP^high^/SSC-A^low^ population), as validated by post-sort imaging (see **Fig. 1**).

Absolute cardiomyocyte counts per ventricle were calculated using the following formula:

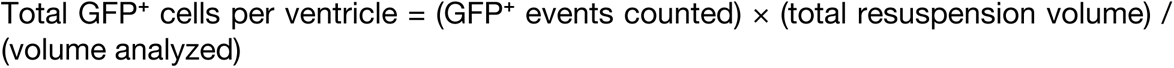

To account for potential variation in fish body size, cardiomyocyte yields were normalized to both resuspension volume and body weight (grams). Each fish was weighed immediately after euthanasia, and normalized yield was calculated as: Normalized yield = (Total GFP⁺ cells per ventricle) / (body weight in grams)

### Experimental Replication and Statistical Analysis

Cardiomyocyte quantification was performed on n=8-10 fish per age group across at least 3 independent experiments conducted on separate days using different fish cohorts. Data are presented as mean ± standard error of the mean (SEM). Statistical comparisons between age groups were performed using one-way ANOVA with Kruskal-Wallis test with Dunn’s multiple comparisons. Statistical significance was set at p < 0.05.

### Flow Cytometric Imaging Using the Attune CytPix

To rigorously assess cardiomyocyte morphology and exclude potential artefacts introduced by fixation, cytospin preparation, or standard flow sorting procedures, we employed the Attune™ CytPix™ Acoustic Focusing Flow Cytometer (Thermo Fisher Scientific) for combined high-throughput flow cytometry and real-time imaging analysis. Six 7-month-old adult male *cmlc2:EGFP* zebrafish were euthanized, and ventricular tissue was dissected and dissociated as described previously to obtain single-cell suspensions. All steps prior to analysis were performed without chemical fixation to ensure assessment of live-cell morphology. Samples were not filtered before loading into the Attune CytPix system. Data acquisition utilized the integrated high-speed brightfield imaging module with a 10x objective, capturing individual cell morphology concurrently with conventional flow cytometry parameters. Live GFP^Hi^/SSC-A^Hi^ cardiomyocytes were identified in the green fluorescence (BL1, 530±15 nm) channel, and size and granularity characteristics were determined using standard forward (FSC-A) and side (SSC-A) scatter profiles. The gating strategy specifically isolated GFP^Hi^/SSC-A^Hi^ events corresponding to ventricular cardiomyocytes (**Fig. 7H**). Cell data were acquired using the Invitrogen™ Attune™ CytPix Flow Cytometer instrument and the Invitrogen™ CytKick™ Max Autosampler at a sample injection rate of 25 µL/minute. For each sample, 50,000 events were collected in “All Events” mode and 10,000 GFP^Hi^/SSC-A^Hi^ cardiomyocytes were imaged in brightfield mode and reviewed using the CytPix software suite (v1.0, Thermo Fisher Scientific). Morphological categories (rounded, rod-shaped, elongated, multipolar) were annotated manually from acquired images, and artefactual features (e.g., membrane blebbing, fragmentation) were excluded from analysis. All analyses were conducted on unfixed cell preparations to ensure accurate representation of physiological morphology. Representative images were exported at full resolution for publication (see **Fig. 7**).

### Cytospin Procedure for Cardiac Cells in Suspension

To prepare cells for cytospin, CytoSep™ Cytology Funnels (including the base holder and fluid chamber) were assembled for use with the Sakura Cyto-Tek® Cytocentrifuge (Model 4323, Sakura). A white filter (M963FW, Simport) was placed into the funnel. Fixed cells in 1-2% paraformaldehyde (PFA) suspension were then centrifuged at 2,000 rpm for 3 minutes onto Epredia™ SuperFrost Plus™ Adhesion slides (12312148, Fisher Scientific). After centrifugation, a hydrophobic barrier was drawn around the sample using a PAP pen (ab2601, Abcam) to prepare the slide for staining. Typically, 100-200 µL of cell suspension was used per sample.

### Immunofluorescence and Antibody Staining

Cells on cytospin slides were washed with 1X PBS, then permeabilized in 1X saponin-based permeabilization/wash buffer (0.2% (w/v) saponin, 2% (v/v) NCS, 1% (w/v) BSA, and 0.02% (v/v) sodium azide in PBS) for 15 minutes. After permeabilization, the buffer was removed, and samples were incubated overnight at 4°C with primary or conjugated antibodies prepared in 1X permeabilization/wash buffer. The next day, the samples were washed three times with 1X PBS before adding secondary antibodies, also prepared in 1X permeabilization/wash buffer, and incubating at room temperature (RT), covered, for 60–120 minutes. In some cases, Wheat Germ Agglutinin, CF555 Conjugate (29076, Biotium), was added and incubated at RT for 30 minutes (1:250 dilution in permeabilization/wash buffer). After this incubation, samples were washed three times with 1X PBS. Finally, a 1:5000 dilution of Vybrant™ DyeCycle Violet in 1X permeabilization/wash buffer was prepared, and samples were incubated with this solution and stored at 4°C until imaging.

Conjugated and primary antibodies used in this study include anti-Cardiac Troponin T (cTnT) Mouse Monoclonal Antibody (13-11, 1:500; ThermoFisher Scientific); Alexa Fluor® 647 Mouse anti-Cardiac Troponin T (cTnT), (1:250; BD Biosciences), PE Mouse anti-Cardiac Troponin T (cTnT), (1:250; BD Biosciences), anti-α-Actinin (Sarcomeric) Vio® R667, REAfinity™ (1:200; Miltenyi Biotec), anti-TTN (Titin) monoclonal antibody (M07), clone 2F12 (mouse, 1:250, Abnova), anti-Myomesin (mouse, mMaC myomesin B4, 1:200, DSHB).

Secondary antibodies (1:500; Thermo Fisher Scientific) used in this study include Alexa Fluor 488 donkey anti-mouse IgG (H + L), and Alexa Fluor 555 donkey anti-mouse IgG (H + L). CF®555 WGA (29076, Biotium, prepared in permeabilization/wash buffer) was added to each slide and incubated covered at room temperature for 30 min. The cells were then washed three times in PBS. Details about the antibodies used in this study can be found in the Supplementary Materials (**key resources table**).

### Confocal Laser Scanning Microscopy

Confocal laser scanning microscopy of cytospun cells and cryosections was performed using a Zeiss LSM900 inverted confocal laser scanning microscope, which includes an upright Zeiss Axio Observer 7, Colibri 5 solid-state LED fluorescence light sources (solid-state laser lines: 405, 488, 561, 640), two Gallium Arsenide Phosphide photomultiplier tubes (GaAsP-PMT), and a motorized stage controlled by ZEN blue 3.4 software. Regular or tile images were acquired using objectives ranging from ×10 (0.45 NA with a WD 2.0, Air Plan-APO UV-VIS–NIR), ×20 (0.8 NA with a WD 0.55, Air Plan-APO UV-VIS–NIR), ×40 (1.3 NA with a WD 0.21, Oil Plan-APO DIC-UV-VIS–IR), and ×63 (1.2 NA with a WD 0.19, Oil Plan-APO DIC-UV-VIS–IR). The ×40 and ×63 objectives were used with immersion oil ImmersolTM 518 F (433802-9010-000, Zeiss) at a 1024 × 1024 or 2048 × 2048 pixels dimension. For z-stack 3D imaging, z-step sizes ranged from 0.25-1 μm, with images acquired under confocal settings using a motorized focus drive. Laser power during imaging was kept below 1%.

### Cardiac resection in adult zebrafish for FACS analysis of *gata4*^+^ cardiac progenitors

*Tg(-14.8gata4:GFP)^ae^*^1^ zebrafish^52^ were maintained under standard conditions at 28°C on a 14-hour light/10-hour dark cycle. Ventricular resections were performed in adults, aged 6 months. Briefly, each fish was anesthetized in 0.016% tricaine solution until opercular movement ceased, then positioned ventral side up on a damp sponge. A small incision was made through the skin and pericardial sac to expose the beating heart, and ∼20% of the ventricular apex was removed using fine iridectomy scissors. Seven days after surgery, zebrafish were euthanized in an overdose of tricaine, and their hearts were immediately excised and immersed in ice-cold PBS. The ventricle was then carefully dissected away from the bulbous arteriosus and the atrium, opened halfway and processed for single-cell dissociation according to the protocols described above.

### Sustained Zebrafish Swimming Exercise Protocol

Adult (7 months) male zebrafish (*Danio rerio*) of the double transgenic line *Tg(cmlc2:EGFP;Tg(-5.1cmlc2:DsRed2-nuc))* were used for the swimming exercise study. All procedures were conducted at 28°C. Prior to the initiation of the exercise regimen, critical swimming speed (Ucrit) was determined individually for each fish. Ucrit testing was performed using custom software that incrementally increased water velocity by 0.5 cm/s every 4 minutes until the maximum speed of the swim tunnel apparatus was reached. The Ucrit values were used to calculate the target exercise intensity for each fish.

Exercise fish (n=10) underwent structured swimming at 70% of average Ucrit for 6 hours/day, 5 days/week, for 4 weeks in a 5L swim tunnel (SW10050, Loligo Systems; 230V/50Hz). Swimming was monitored using a high-speed camera (VE10380, 4MP USB 3.0, Loligo Systems). During the first week, a ramp-up phase was implemented, beginning at 40% of the average Ucrit and increasing daily by 5–10% until the target intensity of 70% was reached. All treatment groups were fed the same diet once daily, with a pre-measured pellet amount, after exercise—that is, all groups were fed simultaneously. This feeding regimen contrasts with the routine (non-experimental) diet, in which BioCORE husbandry staff feed the fish twice daily with both pellets and artemia, totaling four meals per day. All fish underwent a two-week acclimation period to the experimental environment and feeding schedule prior to the start of any experiment. Non-exercised fish (n=10) maintained under standard tank conditions comprised the control group. Cardiac function was assessed using high-frequency echocardiography, performed at two time points: (1) baseline, prior to Ucrit testing, and (2) endpoint, on the final day of the exercise protocol following the last 4-hour swim session.

### High frequency echocardiography

Adult male zebrafish (8 months old) underwent underwater echocardiography using the Vevo3100^®^ Imaging Station (VisualSonics, Amsterdam, Netherlands) equipped with an MS700D high-frequency transducer. Two-dimensional (B-Mode) imaging, as well as color and pulsed-wave Doppler recordings, were obtained in the optimized long-axis view to evaluate either ventricular or atrial function. Image processing and quantification were performed using VevoLab^TM^ software version 5.7.0 (VisualSonics) by a single operator blinded to genotype. B-Mode long-axis images facilitated measurement of ventricular end-diastolic and end-systolic volumes (EDV and ESV), as well as maximal atrial area (AA), with values normalized to body surface area (BSA). Furthermore, ventricular wall motion was assessed using speckle tracking analysis with VevoStrainTM software, allowing calculation of heart rate, ejection fraction (EF), global longitudinal strain (GLS), and global longitudinal strain rate (GLSR).

### High-Throuput Image Analysis

Image analysis was conducted using Fiji (ImageJ). To begin, automatic thresholding was applied using the Huang method with the "dark" option enabled. The threshold was applied without resetting existing parameters. Measurement settings were then configured to include area (in µm²), mean gray value, perimeter, shape descriptors (circularity and aspect ratio), Feret’s diameter, and elliptical fit parameters. All measurements were redirected to the original stitched image file to ensure accurate spatial and intensity data. Subsequently, the Analyze Particles function was used to identify and quantify objects within the size range of 200 to 3500 µm². Overlay masks were generated to visualize detected particles, individual measurements were displayed, and summary statistics were generated.

### Imaging Data Preprocessing of Control vs Exercise Conditions

Raw morphological features were standardised using z-score normalisation across all cells to account for scale differences across feature dimensions. The two biological batch replicates were normalised separately to account for experimental batch differences, and data was later concatenated together to form a single dataset. Each batch was composed of 4 biological replicates each.

### Imaging Data Dimensionality Reduction

Principal component analysis (PCA) was performed using the scikit-learn package, on the normalised dataset comprising 11 features to reduce dimensionality to one dimension. This selection of a single component was justified by comparative analysis with higher-order PCA applications, which revealed that principal component 2 (PC2) contributed minimal additional information, and principal component 1 (PC1) sufficiently extracted the dominant axis of variation.

### Histograms and KDEs

Kernel density estimates (KDEs) were calculated using Gaussian kernels across 100-point subsets, for smooth density estimation using the scipy.stats and matplotlib packages. KDEs were computed for each condition separately and overlaid to visualise population shifts. The control densities were subtracted from the exercise densities to compute differences pointwise along PC1 and these were plotted to visualise the relative shift in phenotypic enrichment. Trends were plotted across the entire dataset and separately for each technical replicate to assess consistency. One of the 8 replicates displayed inverse trends to all others, suggesting a technical error in labelling or in the experiment itself. Thus, this replicate was removed from downstream data analysis.

### Enriched Subset Definition

To define phenotypic populations, the PC1 value of the X-intercept of the trend line, marking the shift from control to exercise enrichment, was used to subset the population into control-enriched (< x-intercept) and exercise-enriched (> x-intercept) groups.

### Plotting and Visualisation

Data visualisations like violin plots and radar plots were generated using the matplotlib and plotly packages. Violin plots were employed to visualise the distribution of feature values across the condition-enriched subsets. Radar plots were generated to visualise and compare the multivariate phenotypic profiles of the control- and exercise-enriched subsets. All features were normalised prior to plotting.

### Image Statistical Significance

While violin plots were generated using pooled data across replicates, statistical comparisons between subsets were performed at the biological replicate level to enhance statistical power using t-tests from the scipy.stats package. P-values were annotated where significant.

### Arresting Heart in Diastole for Confocal Imaging

For all structural microscopy-based analyses, hearts were arrested in diastole prior to tissue fixation. Following explantation, the beating heart was placed in ‘fish fix buffer’ (FFB, 77 mM sodium phosphate dibasic, 23 mM monosodium phosphate, 0.12 mM CaCl2) and the bulbus arteriosus and any extracardiac vessels or debris were removed. Hearts were then immersed in 4% KCl with 1 mg/mL heparin (H3393, Sigma-Aldrich) until diastole had been observed for 2 min. Following arrest, heart was transferred into a 1.5 mL microfuge tube for 4% PFA-assisted fixation. All reagents made up in FFB unless otherwise specified.

### Tissue Fixation, OCT Embedding and Cryo-sectioning

*cmlc2:EGFP;flk1:DsRed2* male adult (12-month-old) hearts were fixed in 4% PFA and incubated for 1 h at room temperature, rinsed three times (5 min each) in PBS, then cryoprotected in 3% (v/v) sucrose overnight at 4°C. Up to 4 hearts–each orientated with the flat face of the ventricle resting on the bottom of the block–were embedded per block in Tissue-Tek OCT, frozen on dry ice, and stored at –80 °C until sectioning. Serial sections of 6 μm thickness were obtained from frozen OCT blocks using a Leica cryostat machine (CM 1950, Wetzlar, Germany) with chamber/collection temperatures –26/–24 °C and mounted on Superfrost Plus slides (Superfrost Plus, #SF41296SP, ThermoFisher Scientific Pty Ltd, US). Sections spanning the valvular plane were collected for histological immunostaining and confocal imaging.

### Ventricular Cell Preparation and FACS for Global Proteomics

Experiments were conducted using the transgenic zebrafish line TE *Tg(cmlc2:EGFP;flk1:DsRed2)*. All samples were obtained from 11-month-old male zebrafish. A total of 12 ventricles were collected and distributed as 4 ventricles per tube across 3 tubes. Ventricles were isolated and halved using a single-edge stainless steel blade, resulting in 4 halved ventricles per tube. Cell dissociation was performed in 2 mL tubes using ZF3.0 buffer, diluted 1:1 (v/v) with 1% BSA. Each tube received 650 µL of the diluted buffer. Samples were incubated at 28°C for 15 minutes on a thermoregulated rocker set to 900 rpm, followed by 20 gentle handshakes and an additional 15-minute incubation. Subsequently, 200 µL of the diluted buffer was added to each tube, and samples were incubated for a further 5 minutes. Dissociation was halted by adding 1 mL of cold FACS buffer, followed by centrifugation at 500 x *g* for 5 minutes using a swing rotor. Most of the supernatant was removed, cells were gently flicked, and 500 µL of cold FACS buffer was added. Cells were stained with DAPI at 1 µg/mL for 10 min. FACS was performed using the BD FACSAria III equipped with an ND2 filter. A 100 µm nozzle was used to sort cells into 200 µL of 1% BSA in 2 mL tubes. Approximately 37,500 cmlc2:EGFP+/flk1:DsRed2-cardiomyocytes and 90,000 cmlc2:EGFP-/flk1:DsRed2+ endothelial cells were sorted across the three tubes. Sorted cells were pelleted at 1,000g using a swing bucket centrifuge. The supernatant was carefully removed, and 400 µL of PBS1X was added to eliminate residual BSA. Cells were pelleted again at 1,000g, and the supernatant was discarded.

### Cell Lysis and Protein Isolation Global Proteomics

Cells pellets were lysed in RIPA buffer (1X) supplemented with protease and phosphatase inhibitors (MCE, mass-spec compatible, without EDTA). Each tube contained approximately 12,500 cells resuspended in 24 µL of RIPA buffer, equating to ∼520 cells/µL. Lysates were mixed thoroughly and sonicated for 5 seconds at 25% amplitude using Branson Tapered Microtip for 1/2" Sonifier Tapped Horns, 1/8" dia (EW-04715-83, Cole-Palmer) attached to Branson Disruptor Horn for Sonifiers 1/2" Dia Stepped and Tapped (1258032, Cole-Palmer) using the Branson SFX550 Sonifier With 3/4" Horn 550 W 20 KHz 240 VAC. Protein samples were stored at −20°C until further processing.

### Sample Preparation, Instrument Method, and MaxQuant Parameters for LC-MS/MS Global Proteomics

RIPA lysed cardiomyocyte and endothelial cell samples were reduced (5 mM DTT, 37°C, 30 min), alkylated (10 mM IA, RT, 30 min), and digested with Sequencing Grade Modified Trypsin (Promega, Catalog Number V5111) at 37°C overnight. Samples were desalted using SDB-RPS stage-tips prepared with two SDB-RPS disks (Empore, Sigma Cat#66886-U) packed in a 200 μL pipette tip as described previously. Each tip was wetted with 100 μL of 100% acetonitrile, followed by equilibration with 30% methanol/1% TFA solution and 0.1% TFA in water. Each tip was loaded with peptides in 1% TFA. The peptides were washed with 0.1% TFA in water, followed by 1% TFA in isopropanol. To elute, 100 μL of 5% ammonium hydroxide in 80% acetonitrile was added. The eluate was then dried in a SpeedVac. Extracted peptides from each clean-up were reconstituted in 10 μL 0.1% (v/v) formic acid, 0.05% (v/v) heptafluorobutyric acid and 2% acetonitrile in water.

### LC-MS/MS Analysis

Digest peptides were separated by nanoLC using an Ultimate nanoRSLC UPLC and autosampler system (Dionex, Amsterdam, Netherlands). Samples (2.5 µl) were concentrated and desalted onto a micro C18 precolumn (300 µm x 5 mm, Dionex) with H2O:CH3CN (98:2, 0.1% TFA) at 15 µl/min. After a 4 min wash the pre-column was switched (Valco 10 port UPLC valve, Valco, Houston, TX) into line with a fritless nano column (75 µm x ∼25 cm) containing C18AQ media (1.9 µm, 120 Å Dr Maisch, Ammerbuch-Entringen Germany). Peptides were eluted using a linear gradient of H2O:CH3CN (98:2, 0.1% formic acid) to H2O:CH3CN (64:36, 0.1% formic acid) at 200 nl/min over 90 min. High voltage 2000 V was applied to a low volume Titanium union (Valco) and the tip positioned ∼0.5 cm from the heated capillary (T=275°C) of an Orbitrap Fusion Lumos (Thermo Electron, Bremen, Germany) mass spectrometer. Positive ions were generated by electrospray and the Fusion Lumos operated in data dependent acquisition mode (DDA).

### Mass Spectrometer Settings

A survey scan m/z 350-1750 was acquired in the orbitrap (resolution=120,000 at m/z 200, with an accumulation target value of 400,000 ions) and lockmass enabled (m/z 445.12003). Data-dependent tandem MS analysis was performed using a top-speed approach (cycle time of 2s). MS2 spectra were fragmented by HCD (NCE=30) activation mode and the ion-trap was selected as the mass analyzer. The intensity threshold for fragmentation was set to 25,000. A dynamic exclusion of 20 s was applied with a mass tolerance of 10 ppm.

### Data Analysis

Peak lists were generated using Mascot Daemon/Mascot Distiller (Matrix Science, London, England) or Proteome Discoverer (Thermo, v1.4) using default parameters, and submitted to the database search program Mascot (version 3.1, Matrix Science). Search parameters were: Precursor tolerance 4 ppm and product ion tolerances ± 0.5 Da; carbamidomethyl (C) and oxidation (M) specified as variable modifications; enzyme specificity was trypsin with 1 missed cleavage allowed; and the SwissProt database (SProt 19_1_24; 570,420 sequences; 206,321,560 residues) was searched. Results were filtered at p<0.05.

Raw data acquired from LC-MS/MS analysis were processed using MaxQuant software (version 2.6.8.0). Peptide and protein identifications were performed via the Andromeda search engine with default settings. Label-free quantification was performed using the MaxLFQ algorithm. Search parameters included: carbamidomethyl (C) and oxidation (M) as variable modifications; enzyme specificity was trypsin with up to 1 missed cleavage allowed; minimum peptide length of 7 amino acids; peptide and protein FDR cut-offs of 1%. The match between runs (MBR) feature was enabled. UniProt FASTA database with decoy strategy was used following MaxQuant default settings.

### Label-free quantitative (LFQ) proteomics analysis

MaxQuant result output contains proteins groups of which 1084 proteins were reproducibly quantified^99^. Of note, the data were normalized based on the assumption that the majority of proteins do not change between the different conditions. Statistical analysis was performed using an in-house generated R script based on the ProteinGroup.txt file. First, contaminant proteins, reverse sequences and proteins identified “only by site” were filtered out. In addition, proteins that were only identified by a single peptide and proteins not identified/quantified consistently in the same condition were also removed.

Following initial protein identification and quantification from the raw mass spectrometry data, an initial quality control assessment was performed. The total number of unique proteins identified with high confidence (e.g., False Discovery Rate [FDR] < 1%) was calculated for each individual sample. This summarization step was used to verify the technical consistency and depth of proteome coverage across all biological replicates for both the cardiomyocyte (CM) and endothelial (EC) conditions. The protein counts per sample were visualized using a bar chart to ensure no sample failures or significant outliers existed before proceeding to data normalization and further statistical analysis (**Fig. S9**).

The LFQ data was converted to log2 scale, samples were grouped by conditions and missing values were imputed using the ‘Missing Not At Random’ (MNAR) method, which uses random draws from a left-shifted Gaussian distribution of 1.8 StDev (standard deviation) apart with a width of 0.3. Protein-wise linear models combined with empirical Bayes statistics were used for the differential expression analyses. Following normalization, we assessed the impact of missing value imputation on the protein intensity distributions (**Fig. S9**). The distribution of reliably quantified proteins was centered and largely consistent between CM and EC samples. The imputation process introduced a distinct, second distribution of low-abundance values, resulting in a bimodal profile for both conditions. This confirms that missing values, presumed to be from proteins below the detection limit, were successfully imputed, thereby preparing the complete dataset for differential expression analysis. The *limma* package from R Bioconductor was used to generate a list of DEPs for each pair-wise comparison. A cutoff of the *adjusted P-value* of 0.05 (Benjamini-Hochberg method) along with a log2 fold change of 1 was applied to determine significantly DEPs in each pairwise comparison.

### STRING-db Protein–Protein Interaction Analysis and Cytoscape Visualisation

Upregulated DEPs in cardiomyocytes (cluster 2) compared to endothelial cells (cluster 1) were selected for protein–protein interaction (PPI) analysis using the STRING-DB database. *Danio rerio* (NCBI taxonomy ID: 7955) was used as the reference species. Of the 328 input proteins, 322 were successfully mapped in STRING. The interaction network was generated using the full STRING network with the following active interaction sources: text mining, experimental data, curated databases, co-expression, neighbourhood, gene fusion, and co-occurrence. The minimum required interaction score was set to medium confidence (0.400). The resulting network was exported to Cytoscape (version 3.10.3), where proteins were clustered using the “Cluster Network MCL (Markov Clustering)” function with default parameters (granularity = 3). MCL is a graph-based clustering algorithm commonly used in bioinformatics to identify protein complexes or functional modules. The same above-described procedures were then applied to upregulated DEPs in endothelial cells (cluster 1, 477 succesfully mapped in STRING) compared to cardiomyocytes, modifying the MCL parameters to granularity = 2. PPI networks were exported as PDF.

### FACS and Low-Input RNA Isolation and Purification

Experiments were performed using the transgenic zebrafish line TE (*cmlc2:EGFP; flk1:DsRed2*). All samples were collected from 12-month-old male zebrafish. A total of nine ventricles were harvested and evenly distributed across three tubes, with three ventricles per tube. Cells were dissociated into single-cell suspensions as previously described. FACS was then used to isolate viable cells (DAPI negative). Endothelial/endocardial cells and cardiomyocytes were directly sorted into 700 µL of QIAzol Lysis Reagent (Qiagen, 79306). Per tube (n=2, pooling three ventricles per tube), approximately 7,000–10,000 cardiomyocytes and 15,000–30,000 endothelial cells were sorted. A higher number of endothelial cells was used for two reasons: (1) endothelial/endocardial cells typically yield less RNA per cell compared to cardiomyocytes, and (2) they are 2–4 times more abundant in the zebrafish heart. Samples were kept on ice throughout the procedure. Total RNA was purified using the Direct-zol RNA Microprep Kit (Zymo Research, R2060), following the manufacturer’s instructions. Briefly, ∼800 µL of QIAzol lysate containing ∼100 µL of sorted cell suspension was mixed with 800 µL of 100% ethanol. The mixture was loaded onto a Zymo-Spin IC Column in a Collection Tube. To ensure complete RNA recovery, two passes of 800 µL each were performed. DNase I treatment was carried out according to the manufacturer’s protocol. All steps were conducted at room temperature, with centrifugation at 12,000 × g. Finally, total RNA was eluted in 15 µL of DNase/RNase-free water by centrifuging for 2 minutes at 12,000 × g. RNA samples were stored at −70 °C until further processing.

### Automated Electrophoresis and RNA Quality Control

Total RNA quality and quantity were assessed using the High Sensitivity RNA ScreenTape assay on the Agilent TapeStation system at the Australian Genome Research Facility (AGRF). RNA Integrity Numbers (RIN) were consistently above 8.3, with an average RIN of 9, indicating high-quality RNA suitable for downstream total RNA-seq applications.

### Whole-Transcriptome Total RNA Sequencing

Total RNA was extracted and prepared for sequencing using the Illumina Stranded Total RNA Prep with Ribo-Zero Plus kit, following the manufacturer’s standard protocols. This library preparation method depletes both ribosomal RNA (rRNA) and globin mRNA to enable a comprehensive, whole-transcriptome analysis of coding and non-coding RNA species. To ensure comparability across samples, all libraries were normalized to a concentration of 21 ng. The low-input libraries were then amplified using 15 cycles of PCR. Sequencing was performed on an Illumina NovaSeq X platform, generating 150 bp paired-end reads with a depth of 50 million read pairs per sample. Primary data processing, including image analysis and real-time base calling, was conducted on the instrument using the NovaSeq Control Software (NCS) v1.3.1.59007 and Real Time Analysis (RTA) v4.29.3. The subsequent conversion of Base Call (BCL) files to FASTQ format was performed using the Illumina DRAGEN BCL Convert pipeline (v07.031.732.4.3.6). All sequencing data met the quality control standards set by the Australian Genome Research Facility (AGRF).

### Data Availability and Bioinformatic Processing

#### Data Storage, Integrity and Read Pre-processing and Quality Control

Raw sequencing data (FASTQ files) were deposited and processed on the National Computational Infrastructure (NCI) Gadi supercomputer under project ID el4. The integrity of all FASTQ files was verified upon transfer using MD5 checksums to ensure no data corruption occurred. A custom script (Trim.sh) was used to perform adapter trimming and remove low-quality bases from the raw paired-end reads. This step ensures that technical sequences from the Illumina sequencing process do not interfere with downstream alignment and analysis.

### Genome Alignment

The quality-filtered reads were then aligned to the Zebrafish reference genome (GRCz11/Zv11) using the STAR (Spliced Transcripts Alignment to a Reference) aligner (v2.7.x or similar). A two-pass alignment strategy was employed to maximize the accuracy of splice junction detection. In the first pass, splice junctions were identified from the initial alignment. This junction information was then used to inform a more sensitive and accurate second alignment pass, generating the final BAM files.

### Post-Alignment Processing and Quality Assessment

Following alignment, a series of post-processing and quality control steps were executed. Custom scripts (STARPostProcess.sh) were used to sort and index the resulting BAM files. To comprehensively assess the quality of the sequencing and alignment, two primary tools were used:

Picard Tools: Alignment metrics, such as mapping rates, duplication levels, and insert size distribution, were calculated using Picard. The output was subsequently plotted in R for visual inspection of data quality across all samples. The RNA-SeQC tool was run on each BAM file to generate a broad set of quality metrics, including gene body coverage, transcript diversity, and strand specificity, further ensuring the reliability of the dataset. The final, processed BAM files and associated quality control reports were used for all subsequent downstream analyses, including gene expression quantification. Post-alignment quality control was performed using *SeqMonk* to generate QC plots and assess data quality. The analysis confirmed the high quality of the sequencing data, with all samples demonstrating high mapping rates to the GRCz11 reference genome and the majority of reads aligning to exonic regions. A high percentage of known genes were detected consistently across all libraries, indicating sufficient sequencing depth for a comprehensive transcriptomic analysis. No samples were excluded from analysis after quality control.

### Bioinformatic Analysis

Initial data filtering was performed on the count matrix to remove genes with low expression. Genes were retained for further analysis only if they exhibited a counts per million (CPM) of at least 1 in one or more sample libraries. For visualization techniques such as clustering and Principal Component Analysis (PCA), the filtered count data was transformed. Differential expression analysis was performed using the DESeq2 package. Count data were normalized using DESeq2’s default size-factor method, and transformed using the regularized log (rlog) [or variance stabilizing transformation (VST)] for visualization and downstream analysis. For select downstream applications, counts were converted to counts per million (CPM), and transformed as log₂(CPM + 4), where a pseudocount of 4 was added to avoid taking the logarithm of zero. Missing values in the dataset were imputed using the median expression value of the corresponding gene across all samples.

### Multi-Omics Data Integration and Comparative Analysis

To identify proteins with corresponding upregulated gene expression, we performed a comparative analysis between two distinct cardiomyocyte-specific datasets. The primary datasets consisted of (i) a list of abundant proteins, or DEPs, identified via mass spectrometry-based proteomics of cardiomyocytes, and (ii) a list of upregulated DEGs identified from cardiomyocyte transcriptomics. The same approach was applied to endothelial cell datasets.

### Data Preprocessing and Normalization

Prior to comparison, both lists were programmatically standardized to ensure consistent matching. All protein and gene identifiers, including official gene symbols (e.g., TNNT2), accession numbers (e.g., zgc:158660), and annotated loci (e.g., ttn.1), were converted to a common lowercase format to eliminate case sensitivity. Extraneous whitespace and formatting artifacts were removed from each entry to create clean, uniform lists of identifiers.

### Protein-Gene Set Overlap Algorithm

A multi-stage matching algorithm was implemented to generate a comprehensive list of overlapping entries between the proteomics and transcriptomics datasets. Direct Matching: An initial comparison was performed to identify identifiers with exact, case-insensitive string matches between the two lists. Paralog-Aware Matching: To account for the teleost-specific whole-genome duplication event in zebrafish (*Danio rerio*), the algorithm was expanded to identify orthologous and paralogous relationships. A match was accepted if a base gene symbol from one list corresponded to an annotated paralog in the other. This rule was applied bidirectionally and included common zebrafish paralog nomenclatures, such as: Matching a base symbol to a paralog with an alphabetical suffix (e.g., proteomics tnnt2 matching transcriptomics tnnt2a), or matching between two different alphabetical paralogs of the same gene family (e.g., proteomics myom1b matching transcriptomics myom1a or Myom1), or matching gene symbols with numerical or complex annotations (e.g., proteomics ttn.1 or ttn.2 matching transcriptomics ttn). The same approach was applied to endothelial cell datasets.

### Analysis, Verification, and Tooling

The comparison algorithm was implemented using custom Python scripts. The analysis, execution of scripts, and iterative refinement of the matching logic were facilitated by a large language model (Gemini 2.5 Pro, Google). The process involved multiple passes of computational analysis followed by logical and human-based verification to resolve ambiguities and ensure that the final, consolidated list was both accurate and comprehensive. The final output is a curated list of 284 unique proteins/genes identified as present in both the CM (CM vs ECs) upregulated proteome and the CM (CM vs ECs) upregulated transcriptome. On the other hand, 376 unique proteins/genes were identified in ECs.

### AI-Driven Functional Annotation and Enrichment Analysis of Multi-Omics Integrated Datasets

The functional significance of the final, curated lists of protein-genes overlapping proteins was determined through a knowledge-based analysis performed by a large language model (Gemini, Google). The process began with the input of a standardized list of official gene symbols. The model then executed a multi-step analytical workflow:

#### Functional Annotation

The LLM leveraged its extensive internal knowledge base, which has been trained on a vast corpus of scientific literature and bioinformatics databases (including Gene Ontology, UniProt, and KEGG). Each protein identifier in the input list was systematically mapped to its associated Gene Ontology (GO) terms across the three primary domains: Biological Process (BP), Cellular Component (CC), and Molecular Function (MF).

#### Inferred Enrichment and Thematic Clustering

Following comprehensive annotation, the model performed an inferred enrichment analysis. This involved identifying and quantifying the frequency of all GO terms associated with the input list. The terms were then thematically clustered and ranked based on the number of genes from the list that mapped to them. This method identifies the most dominant functional signatures and cellular localizations represented within the dataset.

#### Statistical Representation and Data Synthesis

To represent the statistical significance of these themes, the most prominent GO terms were presented in summary tables. These tables included the number of genes from the input list associated with each term (Count) and representative protein examples. Key metrics like Fold Enrichment and P-value were generated based on the model’s analysis of the input list’s composition against its general knowledge of protein function distribution, simulating the output of standard enrichment tools for interpretability.

### Statistical Analysis

For statistical analysis, unless otherwise specified, all results obtained from independent experiments are reported as means ± standard errors of means (SEM) of multiple replicates. Comparisons between two groups of normally distributed and not connected data were performed using unpaired, nonparametric Student’s t-test. For independent samples, the Mann-Whitney U test was used. Multiple group comparisons were performed by one-way analyses of variance analyses (ANOVA, for one independent variable) or two-way ANOVA (for two independent variables), followed by Tukey’s post hoc comparison (GraphPad Prism version 10, La Jolla, CA). For one-way ANOVA we performed Kruskal-Wallis test with Dunn’s multiple comparisons. Unless otherwise indicated, “*n*” in Figure Legends represents the number of animals or independent biological samples or replicates per group used in the indicated experiments. P-values < 0.05 were considered statistically significant. \**p* < 0.05, \*\**p* < 0.01, \*\*\**p* < 0.001, \*\*\*\**p* < 0.0001.

## AUTHOR CONTRIBUTIONS

Conceptualization: O.C.

Methodology: O.C., G.D.S., C.S., M.D., C.T., R.C., L.Z., A.G.R.

Validation: O.C., G.D.S., C.S., M.D., C.T., L.Z., J.G.L.

Formal analysis: O.C, G.D.S., C.S., M.D., L.Z.

Investigation: O.C., G.D.S., C.S., M.D., R.C., A.G.R., J.G.L.

Resources: O.C., R.P.H., D.F., E.W., J.G.L.

Supervision: O.C.

Data curation: O.C., G.D.S., C.S., M.D., C.T. L.Z.

Project administration: O.C.

Funding Acquisition: O.C., R.P.H., D.F., E.W.

Writing–original draft: O.C.

Writing–review & editing: All authors.

## Competing interests

The authors declare that they have no competing interests.

**Correspondence** and requests for materials should be addressed to lead contact, Osvaldo Contreras.

## Acknowledgments

We are grateful to Kelly Smith, Jamie Vandenberg, and Renee Chow for feedback on the project, to Dashika Palipana for discussion of the data, and Jasmina Cvetkovska for blinded analysis of zebrafish echography datasets. We also thank Paul Young and David Humphreys for RNA-seq quality control, Cecelia Jenkin and Aaron Hay for zebrafish husbandry (VCCRI BioCore), and Bernice Stewart for her administrative contributions. Thanks to Arjun Verma, Christopher Augood, and Naushad Moti from Thermo Fisher Scientific for facilitating the Attune CytPix, and the Victor Chang Cardiac Research Institute Innovation Centre (funded by the New South Wales Government Ministry of Health), GWCCG Core Facility Flow (Eric Lam and Yasmin Husaini) and Histopathology Facilities (Anaïs Zaratzian and Andrew Da Silva) at Garvan Institute of Medical Research (GIMR) for their infrastructure support and assistance. Figures were created with Adobe Illustrator and Adobe Photoshop 2024, Adobe.

## Data and materials availability

All data needed to evaluate the conclusions in the paper are present in the paper and/or the Supplementary Materials. Prior to publication we will deposit raw and processed sequencing data and proteomics datasets in appropriate public repositories and provide accession numbers in the manuscript.

