## Supplemental Figure 1 for "Proteo-transcriptomics and morphometrics of teleost cardiac cells define regulatory networks and exercise-induced cardiomyocyte hypertrophy and hyperplasia"

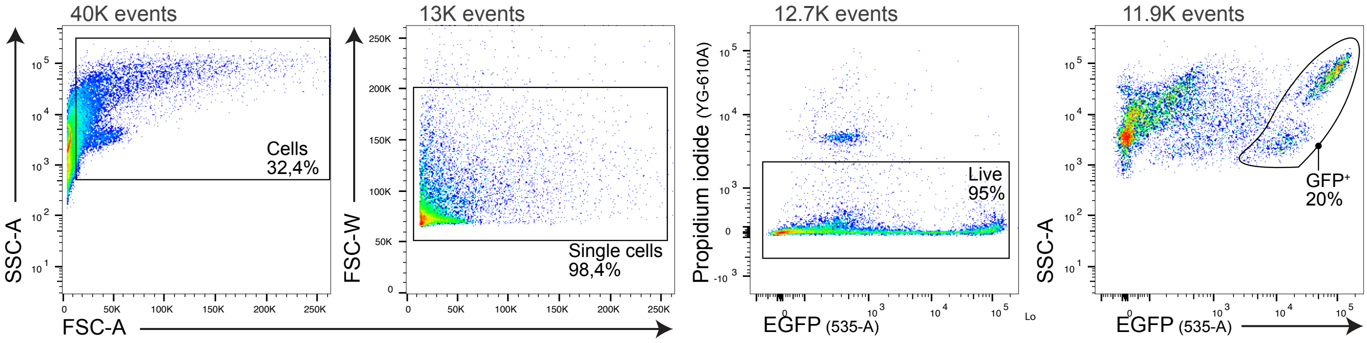


**Fig S1. Flow cytometry strategy for identifying live ventricular cardiomyocytes in cmlc2:EGFP zebrafish.**

(A) Representative gating strategy following enzymatic dissociation of adult ventricular tissue. Initial gates were applied to total events (40,000; 40K), followed by selection of live cells based on exclusion of Zombie Yellow, propidium iodide, and DAPI. Final gating was performed on side scatter area (SSC-A) versus EGFP fluorescence to identify cmlc2:EGFP-positive cardiomyocytes. Data shown are from 7-month-old male zebrafish (N = 4), with a total of 8 ventricles analyzed (2 ventricles per tube).
