## Supplemental Figure 2 for "Proteo-transcriptomics and morphometrics of teleost cardiac cells define regulatory networks and exercise-induced cardiomyocyte hypertrophy and hyperplasia"

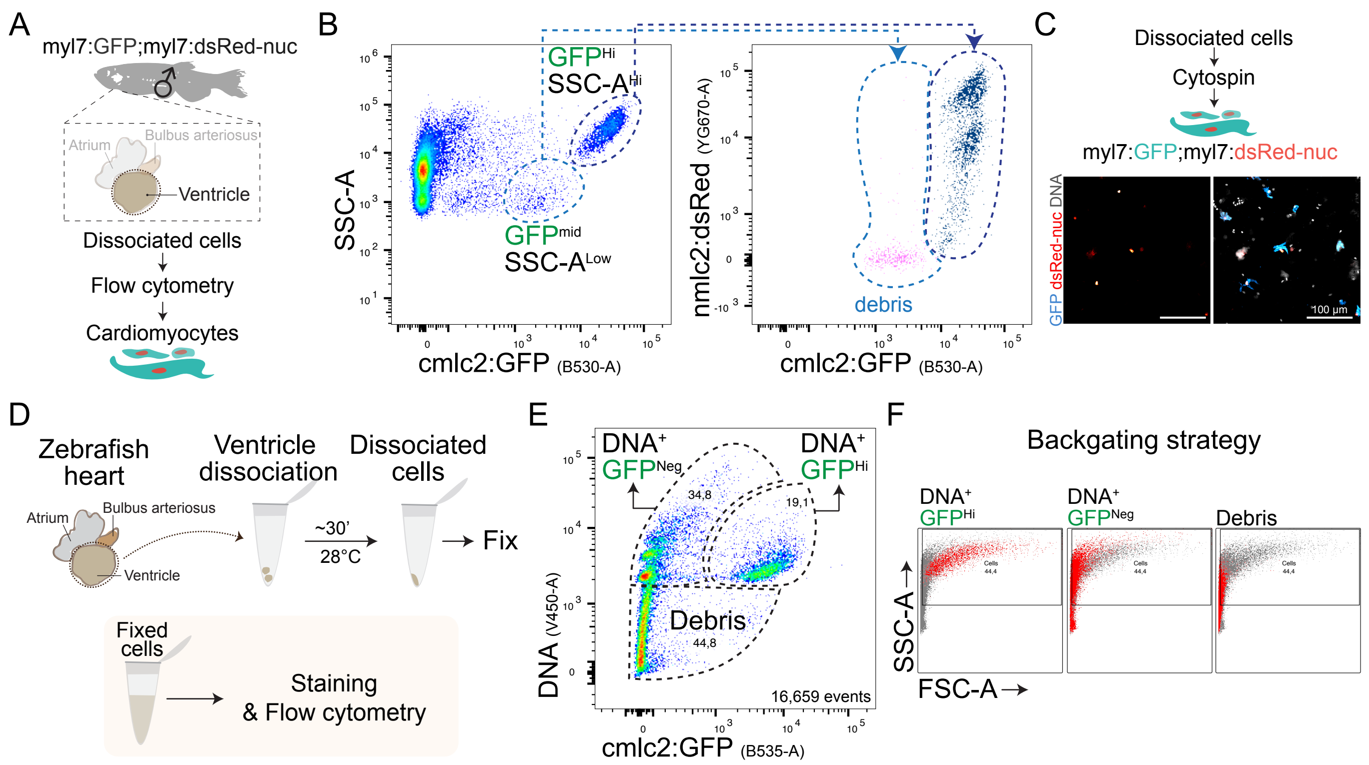


**Fig. S2. Characterization of ventricular cardiomyocytes in zebrafish via flow cytometry and imaging.**
(A) Schematic of fish ventricle isolation, dissociation, and flow cytometry using double transgenic myl7:GFP;myl7:dsRed-nuc adult male fish. (B) Post-dissociation flow cytometry analysis of zebrafish ventricular live cells, displaying dsRed and GFP double-positive ventricular cardiomyocytes based on SSC-A and GFP fluorescence intensity gating strategy. (C) Representative laser confocal images of dsRed and GFP cardiomyocytes. (D) Schematic of ventricle isolation, dissociation, PFA-fixation, and flow cytometry. (E) Post-fixation flow cytometry analysis of dissociated cells, showing three distinct clusters based on DNA and GFP fluorescence intensity. (F) Backgating strategy of the three clusters identified in (E), visualized using SSC-A vs. FSC-A.
