## Supplemental Figure 3 for "Proteo-transcriptomics and morphometrics of teleost cardiac cells define regulatory networks and exercise-induced cardiomyocyte hypertrophy and hyperplasia"

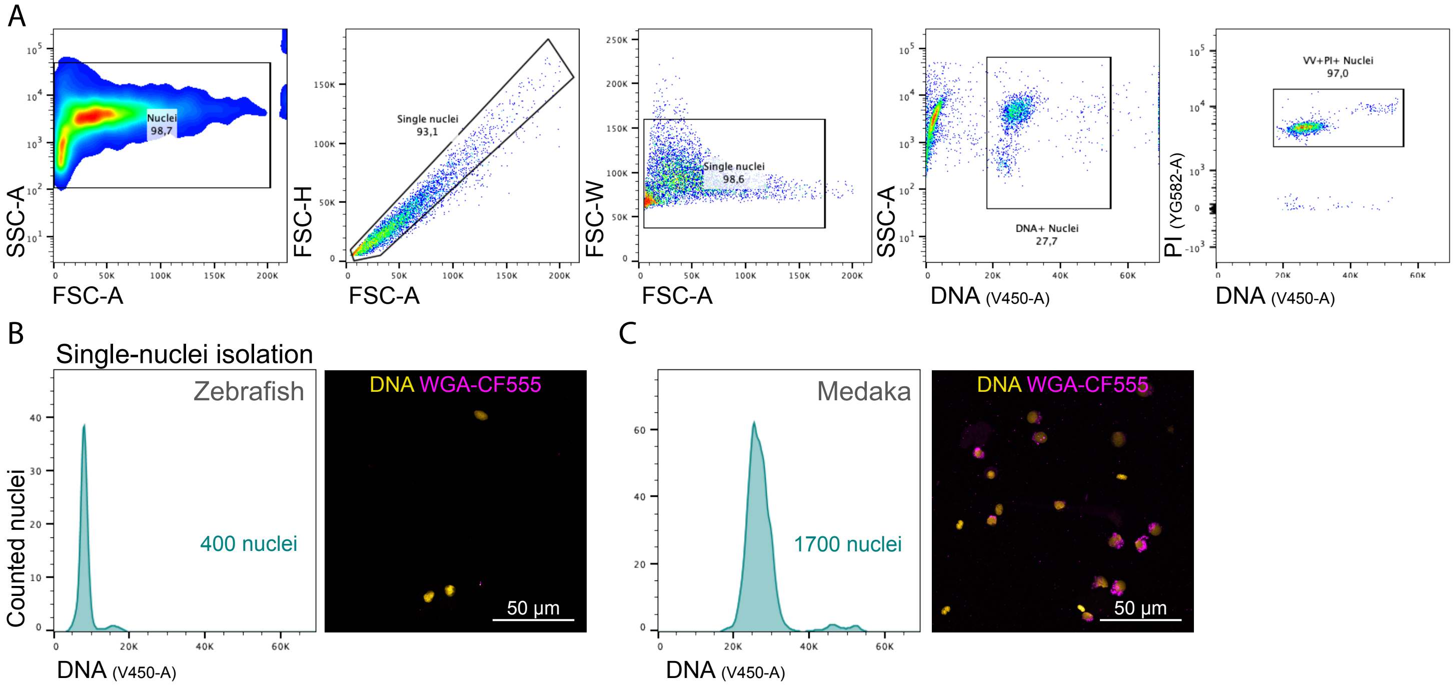


**Fig. S3. Flow cytometric and imaging analysis of single-nuclei isolated from dissociated ventricular tissue.**
(A) Representative flow cytometry plots showing the identification of DNA-stained single nuclei based on side scatter area (SSC-A) following ventricular dissociation. (B, C) Comparative analysis of single-nuclei from zebrafish (B) and medaka (C), including flow cytometry histograms of relative DNA fluorescence intensity. Confocal laser microscopy images confirm nuclear identity, with WGA-CF555 was used to stain cell membranes, ER, and the nuclear envelope.
