## Supplemental Figure 4 for "Proteo-transcriptomics and morphometrics of teleost cardiac cells define regulatory networks and exercise-induced cardiomyocyte hypertrophy and hyperplasia"

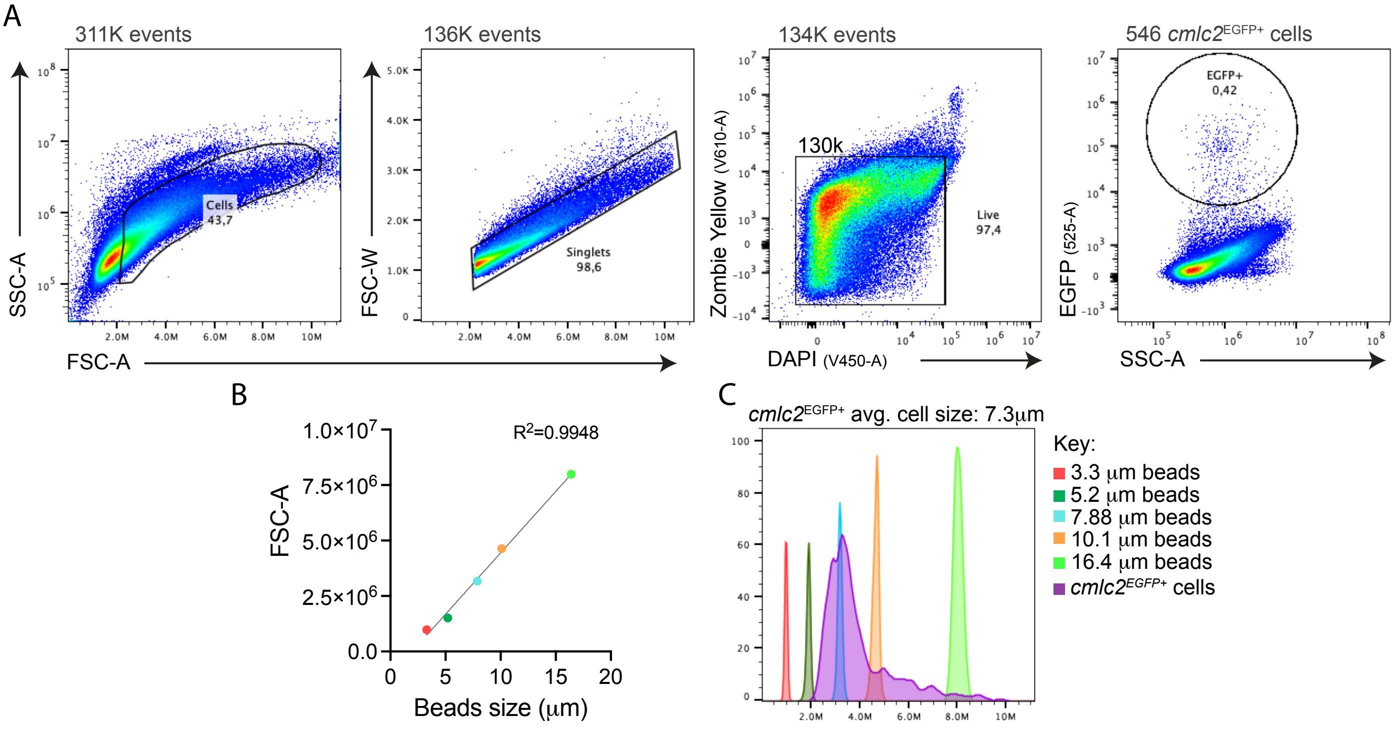


**Fig. S4. Single-cell dissociation and flow cytometric analysis of embryonic cmlc2:EGFP⁺ zebrafish cardiomyocytes.** (A) Representative flow cytometry plots showing the gating strategy for live cells (Zombie Yellow⁻/DAPI⁻) from 28.5 hpf zebrafish embryos following enzymatic dissociation. cmlc2:EGFP⁺ cardiomyocytes are highlighted within the live cell population. (B) Correlation between forward scatter area (FSC-A) and the diameter of calibration beads of known sizes. (C) Quantification of embryonic EGFP⁺ cardiomyocyte size distribution based on FSC-A values, benchmarked against calibration beads shown in (B).
