## Supplemental Figure 5 for "Proteo-transcriptomics and morphometrics of teleost cardiac cells define regulatory networks and exercise-induced cardiomyocyte hypertrophy and hyperplasia"

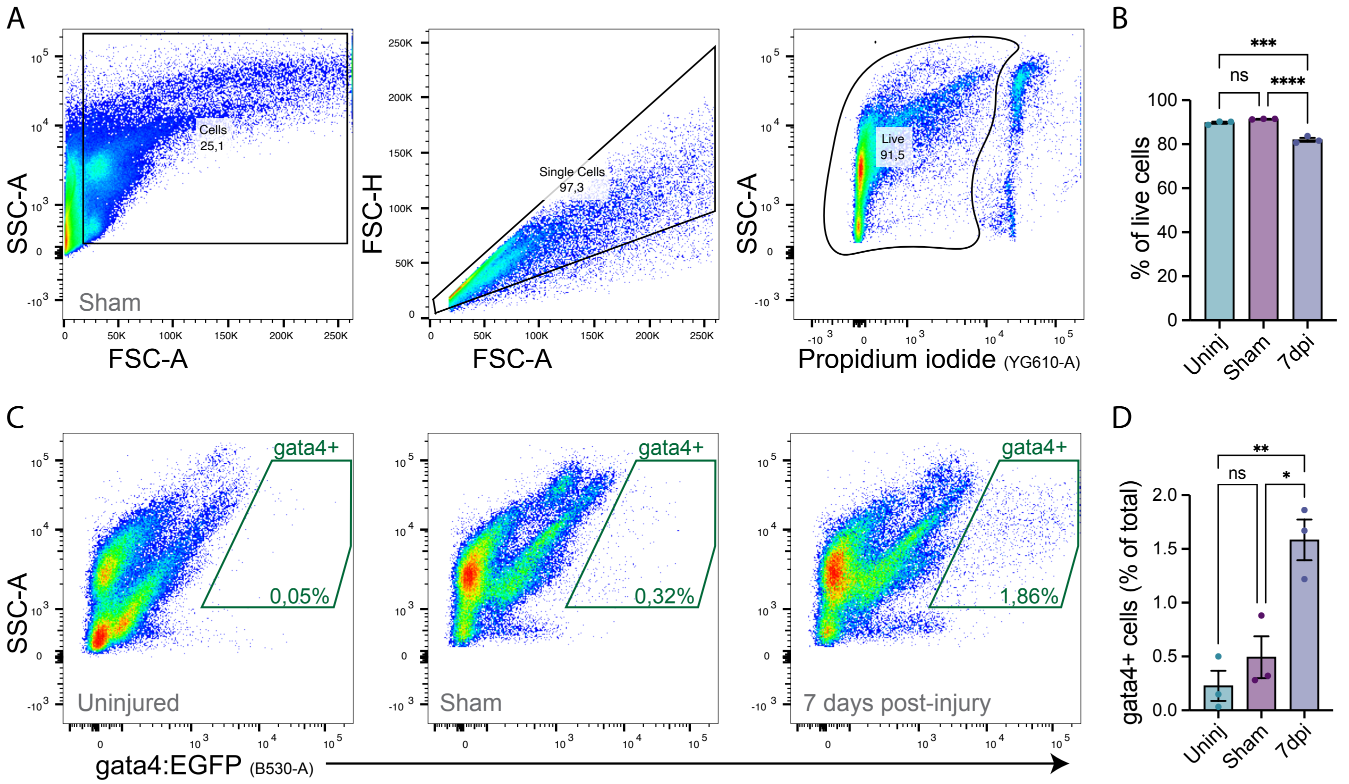


**Fig. S5. Fluorescence-activated cell sorting (FACS) analysis of regenerating ventricular gata4+ cardiomyocyte progenitors in adult zebrafish.**

(A) Representative flow cytometry gating strategy to characterize gata4+ cells based on SSC-A and propidium iodide intensity to characterize viable cells.

(B) Percentage of viable (PI–) cells recovered from ventricular cell suspensions in uninjured controls (Uninj), sham-operated hearts (Sham) and 7 days post injury (7 dpi). Data are mean ± SEM; n = 3 (n= 2, pooled ventricles) per group; one-way ANOVA with Tukey's post hoc test.

(C) Representative EGFP fluorescence histograms of the PI– population in each experimental condition, highlighting the gata4+ (EGFP+) cluster, showing an increase in the number of gata4+ cardiac progenitor cells at 7dpi.

(D) Quantification of gata4+ (EGFP+) cells as a proportion of total viable cells (P3) for Uninj, Sham and 7 dpi samples. Data are mean ± SEM; n = 3 per group; * $p < 0.05$ versus Sham, ** $p < 0.01$ versus Uninj.; one-way ANOVA with Tukey's post hoc test.
