## Supplemental Figure 6 for "Proteo-transcriptomics and morphometrics of teleost cardiac cells define regulatory networks and exercise-induced cardiomyocyte hypertrophy and hyperplasia"

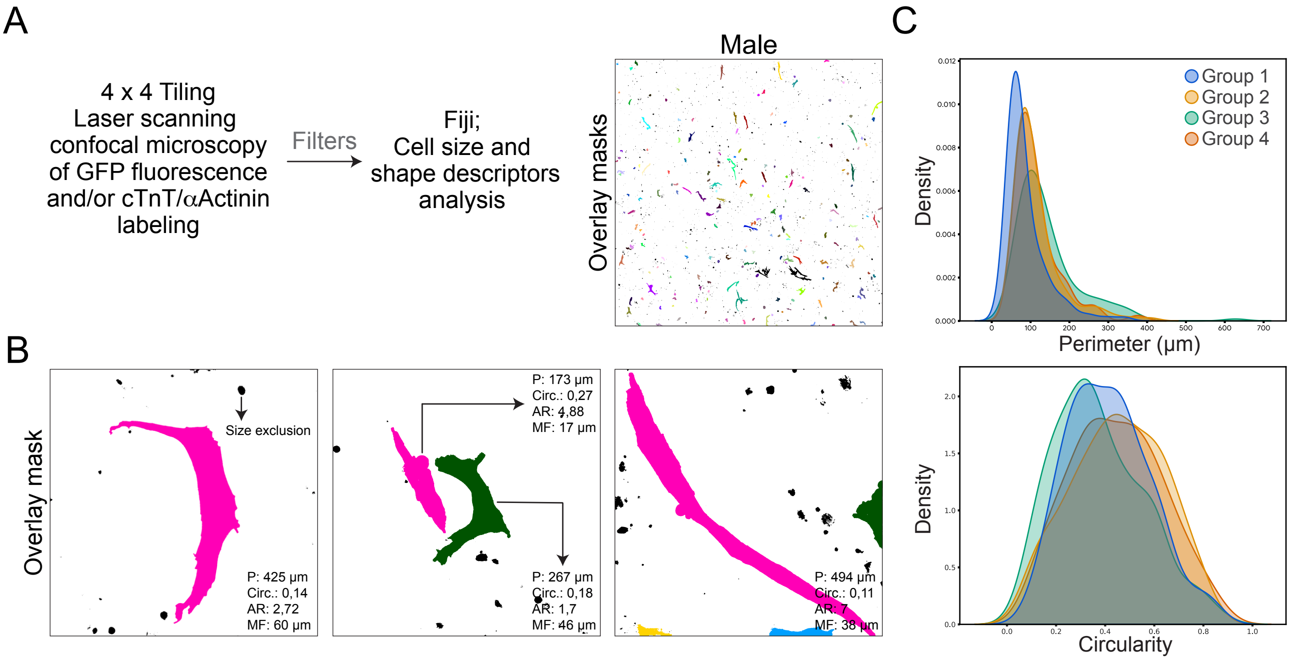


**Fig. S6. Morphological analysis of cmlc2:GFP+ cardiomyocytes from adult male zebrafish ventricles using shape descriptors.**

(A) Representative laser scanning confocal microscopy images (4×4 tile scans) of ventricular cardiomyocytes from a 7-month-old adult male zebrafish (N = 8), visualized via GFP fluorescence and/or immunostaining for cTnT and α-actinin. Overlay masks were applied to identify individual cells. Cells or cell clusters shown in black did not meet the perimeter threshold and were excluded from further analysis. (B) Example ROI overlay masks were generated for individual cells and analyzed using Fiji software to extract shape descriptors. Morphological parameters included perimeter (P), circularity (Circ.), aspect ratio (AR), and minimum Feret diameter (MF). (C) Quantitative comparisons of perimeter and circularity descriptors are shown across four biological groups (each group representing N = 2), revealing heterogeneity in cardiomyocyte size and shape.
