## Supplemental Figure 7 for "Proteo-transcriptomics and morphometrics of teleost cardiac cells define regulatory networks and exercise-induced cardiomyocyte hypertrophy and hyperplasia"

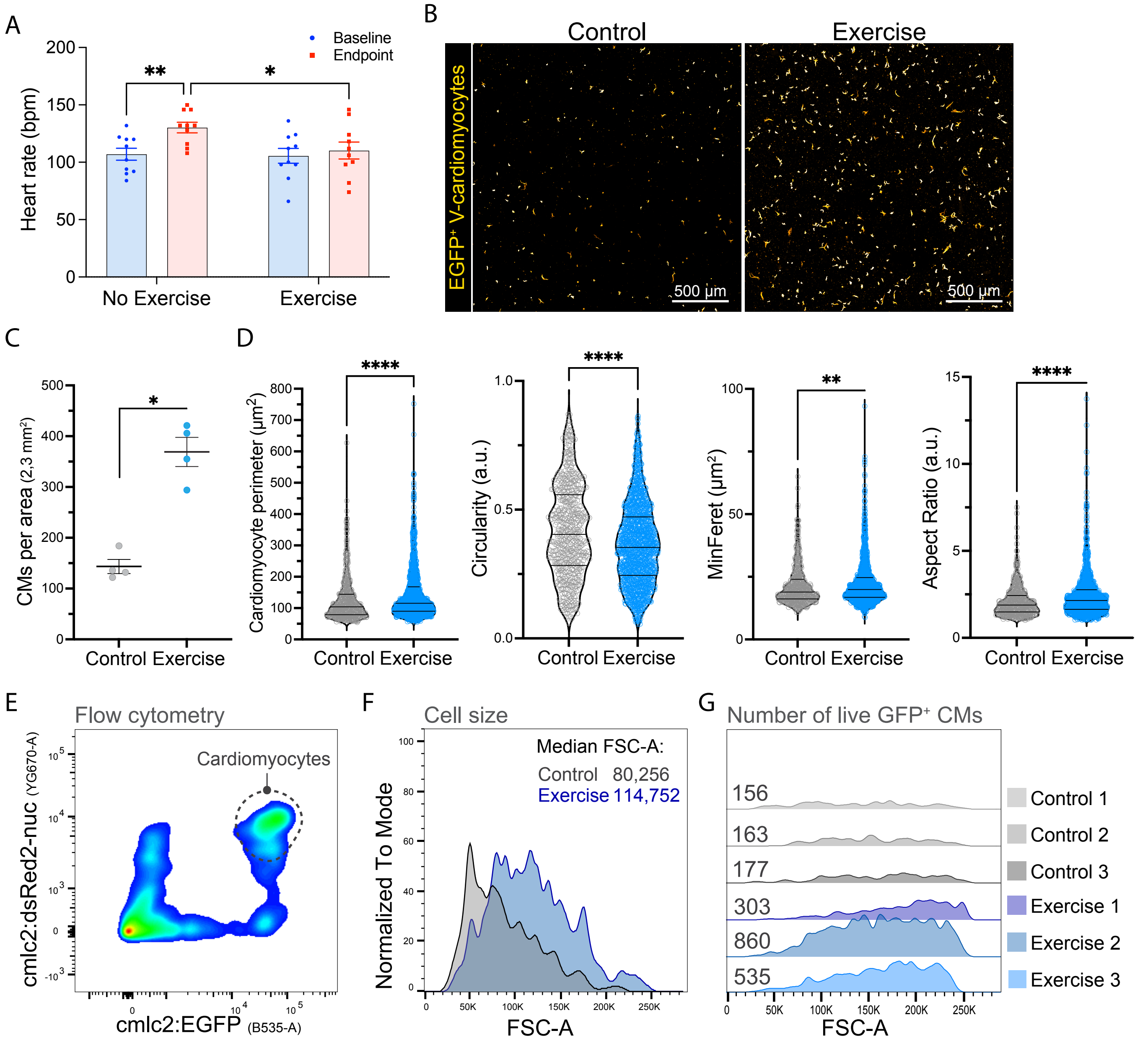


**Fig. S7. Swimming exercise induces ventricular cardiomyocyte hypertrophy and hyperplasia.**
(A) Heart rate (HR) after a 4-week protocol in exercised and non-exercised WT fish. Baseline (blue bars) and endpoint measurements (red bars) are shown. Data are expressed as mean ± SEM. Two-way ANOVA with post hoc Fisher's LSD: *P<0.05; **P<0.01, n = 10. (B) Representative laser confocal images of GFP+ cardiomyocytes. GFP fluorescence is shown in orange hot. (C) Quantification of the number of cardiomyocytes per area. *** P<0.05; nonparametric Mann-Whitney test, n = 4, totaling 10 male ventricles. (D) Quantification of cardiomyocyte perimeter, circularity, MinFeret diameter, and aspect ratio (AR). ****P<0.0001; **P<0.005; nonparametric Mann-Whitney test. (E) Post-dissociation flow cytometry detection of EGFP and nuclear DsRed2-labeled cardiomyocytes using cmlc2 double-transgenic fish. (F) Representative flow cytometry plots illustrating FSC-A (cell size) of cardiomyocytes (n = 3). (G) Representative flow cytometry plots of FSC-A (cell size), showing the number of live cardiomyocytes using equal flow cytometric volume per tube (n = 3).
