## Supplemental Figure 8 for "Proteo-transcriptomics and morphometrics of teleost cardiac cells define regulatory networks and exercise-induced cardiomyocyte hypertrophy and hyperplasia"

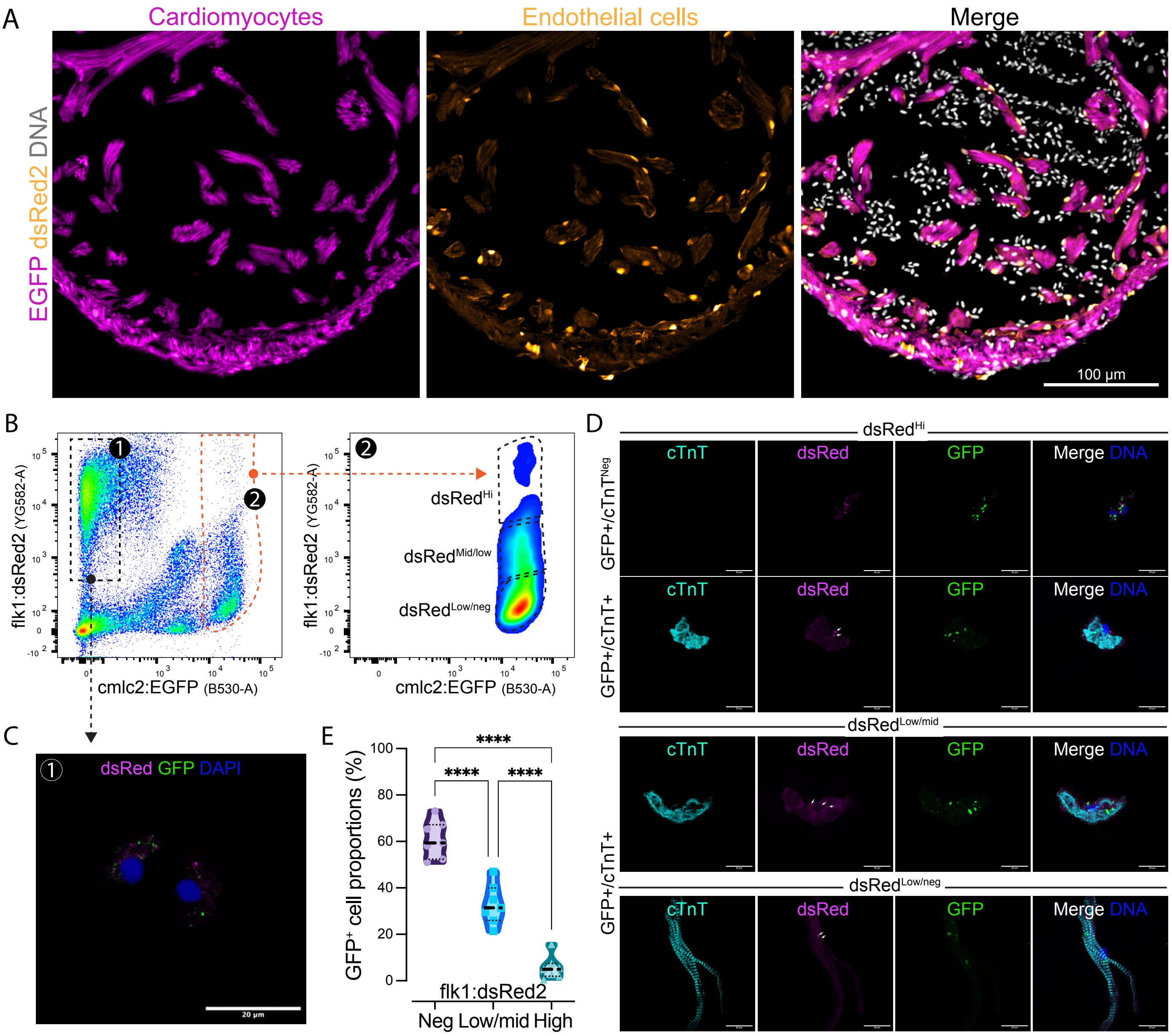


**Fig. S8. Confocal imaging, flow cytometric characterisation, and FACS-based isolation of endothelial and cardiomyocyte populations from cmlc2:EGFP; flk1:dsRed2 double-transgenic zebrafish.**
(A) Z-stack confocal imaging of EGFP and dsRed2 fluorescence, together with DNA staining, in ventricles of an adult male cmlc2:EGFP; flk1:dsRed2 zebrafish.
(B) Flow cytometry analysis of live ventricular cells from adult zebrafish (6–11 months old), identifying flk1:dsRed2+ endothelial cells (Cluster 1, C1) and cmlc2:EGFP$^{+}$ cardiomyocytes (Cluster 2, C2) based on fluorescence intensity. GFP+ cells (C2) were further subdivided into three dsRed2 fluorescence subpopulations: dsRed$^{High}$, dsRed$^{Low/Mid}$, and dsRed$^{Negative}$, and sorted by FACS.
(C) Representative confocal images of FACS-isolated flk:dsRed2$^{+}$ endothelial cells (C1).
(D) Confocal images of cytospun GFP+ cells from C2, showing distinct dsRed2 fluorescence subpopulations. dsRed2 puncta were observed in non-cardiomyocyte (cTnT+) endothelial cells and in a small subset of cTnT$^{+}$ cardiomyocytes. No significant cell doublets were detected.
(E) Quantification of GFP$^{+}$ cells from C2 across dsRed2 fluorescence categories (Negative, Low/Mid, High). ****P<0.0001 by one-way ANOVA with Kruskal-Wallis's post-test; n=7 (14 ventricles total).
