## Supplemental Figure 9 for "Proteo-transcriptomics and morphometrics of teleost cardiac cells define regulatory networks and exercise-induced cardiomyocyte hypertrophy and hyperplasia"

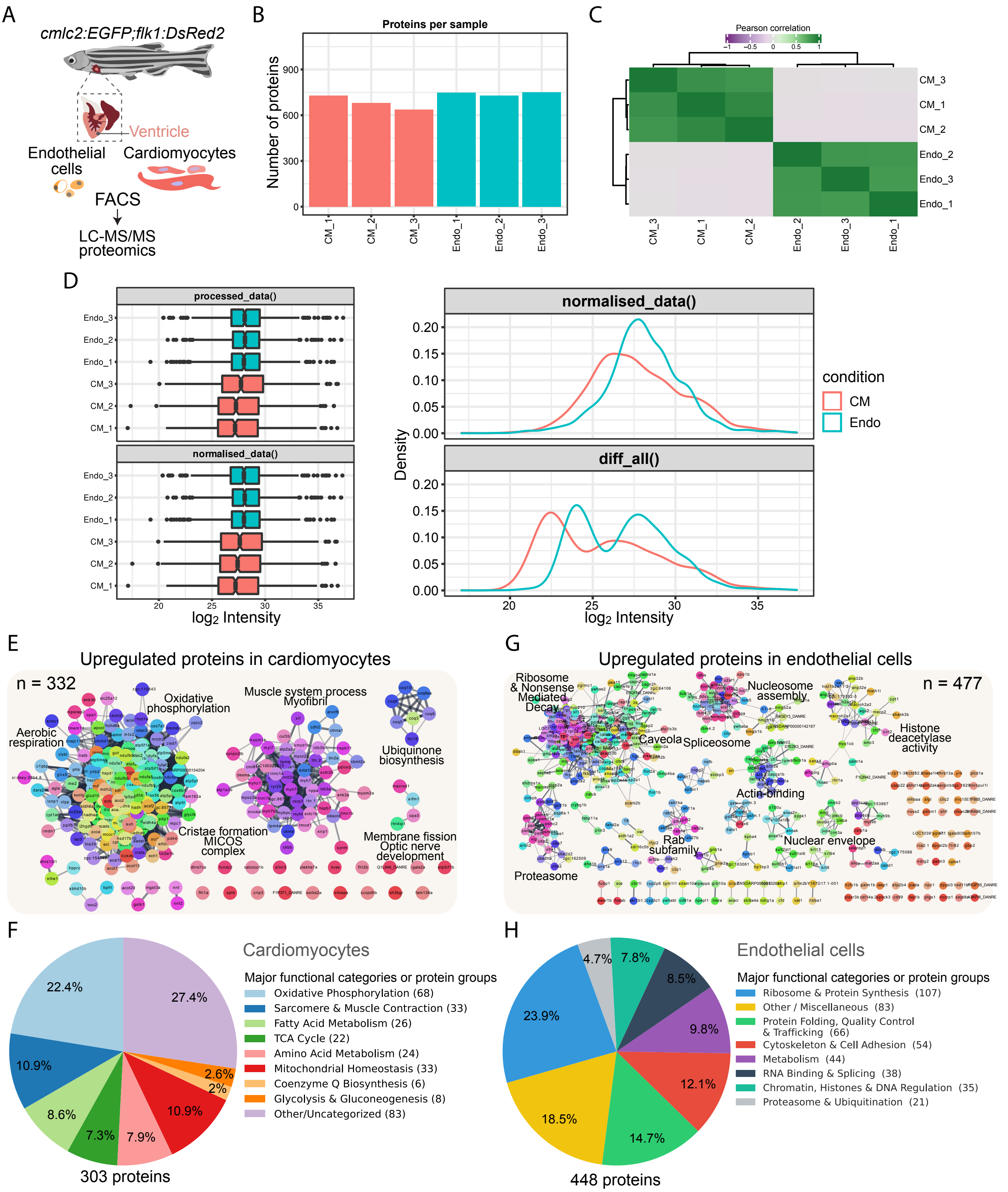


**Fig. S9. Global proteome profiling of ventricular cardiomyocytes and endothelial/endocardial cells.** (A) Schematic of the workflow for isolating dual populations of cardiac cells using cmlc2:EGFP and flk1:DsRed2 transgenes via FACS, followed by protein purification and untargeted proteomics. (B) A sample correlation matrix demonstrating clear separation between cardiomyocyte and endothelial/endocardial cell types. (C) The number of proteins quantified per sample after pre-processing, showing a similar number of detected proteins across all samples. (D) The distribution of missing values and protein expression levels before and after imputation, illustrating the effect of the imputation process. (E) Protein-protein interaction (PPI) network analysis (STRING-db) of differentially expressed proteins (DEPs) enriched in cardiomyocytes (cluster 2). The analysis reveals a highly connected network of proteins associated with aerobic respiration, oxidative phosphorylation (OXPHOS), mitochondrial cristae formation, and cardiac muscle/myofibrils. (F) Pie chart illustrating the proportion of major Gene Ontology (GO) functional categories for proteins enriched in cardiomyocytes. (G) PPI network analysis (STRING-db) of DEPs enriched in endothelial cells (cluster 1). This network shows diverse clusters related to ribosomal biogenesis, protein translation, the proteasome, nucleosome assembly, HDAC activity, actin binding, the nuclear envelope, and the Rab subfamily. (H) Pie chart illustrating the proportion of major GO functional categories for proteins enriched in endothelial cells.
