## Supplemental Figure 10 for "Proteo-transcriptomics and morphometrics of teleost cardiac cells define regulatory networks and exercise-induced cardiomyocyte hypertrophy and hyperplasia"

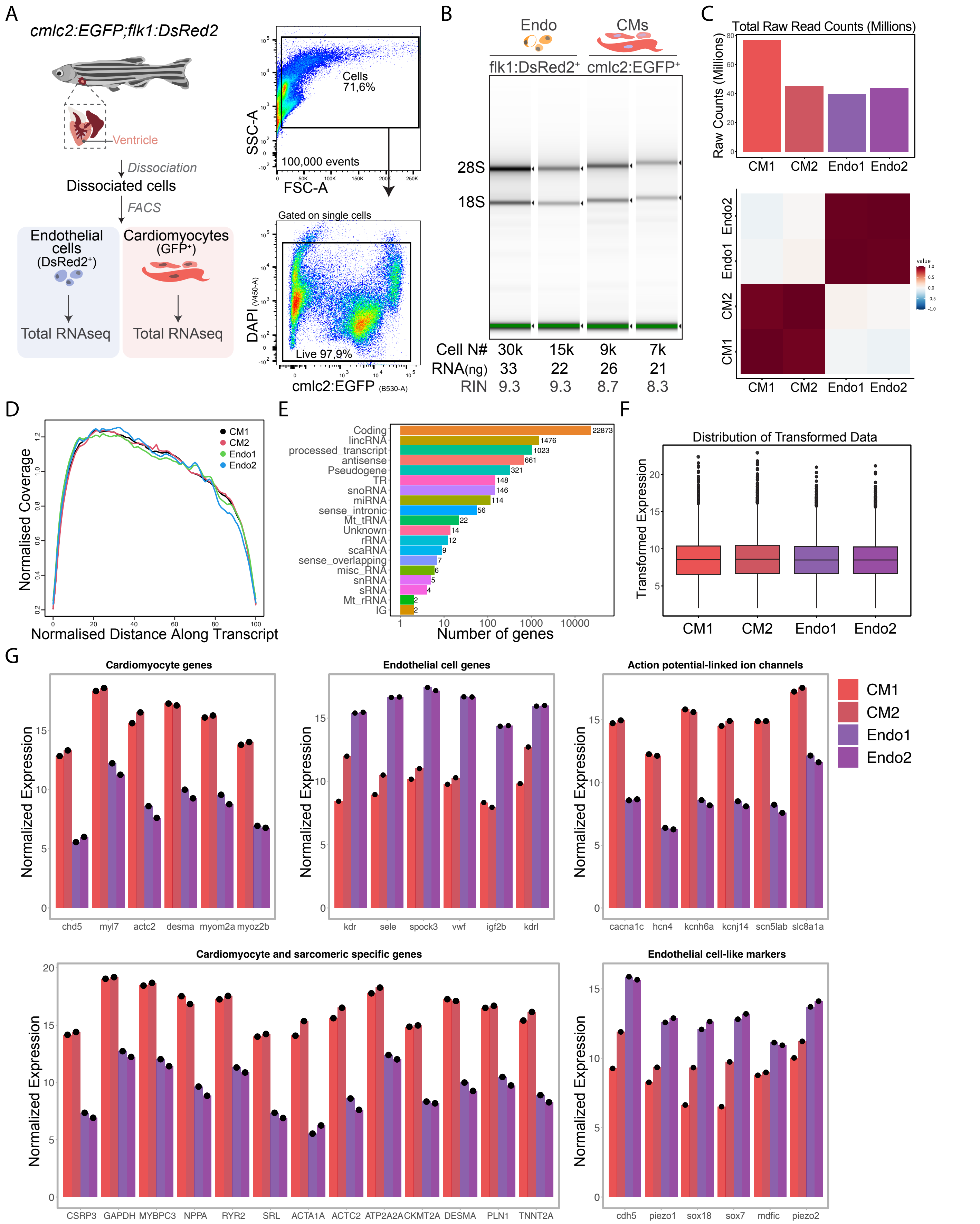


**Fig. S10. Transcriptional profiling of ventricular cardiomyocytes and endothelial/endocardial cells.**
(A)The schematic of the FACS sorting strategy (A) outlines the isolation of CMs (EGFP+) and Endothelial/Endocardial cells (Endo; DsRed2+). cmlc2:EGFP; flk1:DsRed2 zebrafish ventricles for total RNA sequencing. High Total RNA quality assessment (B) is confirmed by RIN≥8.3 across all samples. Total Raw Read Counts (Millions) and the sample correlation heatmap (C) demonstrate high sequencing depth and reproducibility between replicates (CM1/CM2 and Endo1/Endo2), with a clear transcriptional distinction between the two cell types. Data quality checks include the normalized coverage across transcripts (D), which shows uniform read distribution with minimal bias, and the distribution of mapped reads across different RNA biotypes (E), which is dominated by Coding transcripts. The box plots illustrating the distribution of transformed expression data (F) confirm successful global normalization. Finally, the validation of cell-type purity (G) confirms successful isolation by showing the expected differential expression.
